# Functional unit and regulatory mechanisms of supergene in female-limited Batesian mimicry of *Papilio polytes*

**DOI:** 10.1101/2022.02.21.480812

**Authors:** Shinya Komata, Shinichi Yoda, Yûsuke KonDo, Souta Shinozaki, Kouki Tamai, Haruhiko Fujiwara

## Abstract

Supergenes are sets of genes and genetic elements that are inherited like a single gene and control complex adaptive traits, but their functional roles and units are poorly understood. In *Papilio polytes*, female-limited Batesian mimicry is thought to be regulated by a ~130kb inversion region (highly diversified region: HDR) containing three genes, *UXT*, *U3X* and *doublesex* (*dsx*) which switches non-mimetic and mimetic types. To determine the functional unit, we here performed electroporation-mediated RNAi analyses (and further Crispr/Cas9 for *UXT*) of genes within and flanking the HDR in pupal hindwings. We first clarified that non-mimetic *dsx-h* had a function to switch from male to non-mimetic female and only *dsx-H* isoform 3 had an important function in the formation of mimetic traits. Next, we found that *UXT* was involved in making mimetic type pale-yellow spots and adjacent gene *sir2* removed excess red spots in hindwings, both of which refine more elaborate mimicry. Furthermore, downstream gene networks of *dsx, U3X* and *UXT* screened by RNA sequencing showed that *U3X* upregulated *dsx* expression and repressed *UXT* expression. These findings demonstrate that a set of multiple genes, not only inside but also flanking HDR, can function as supergene members, which extends the definition of supergene unit than we considered before. Also, our results indicate that *dsx-H* functions as the switching gene and some other genes such as *UXT* and *sir2* within the supergene unit work as the modifier gene.

**Article summary:** Supergenes are thought to control complex adaptive traits, but their detailed function are poorly understood. In *Papilio polytes*, female-limited Batesian mimicry is regulated by an ~130kb inversion region (highly divergent region: HDR) containing three genes. Our functional analysis showed that *doublesex* switches the mimicry polymorphism, and that an inside gene *UXT* and an outside gene *sir2* to the HDR work to refine more elaborate mimicry. We here succeed in defining the unit of mimicry supergene and some novel modifier genes.

## Introduction

Batesian mimicry is a phenomenon in which a non-toxic species (mimic) escapes predation from predators such as birds by mimicking the appearance, colors, shape, and behavior of a toxic and unpalatable species (model) (Bates 1862) and can be achieved only when multiple traits are properly combined. For example, the butterfly’s wing pattern is composed of various colors and complex patterns, and unless almost all wing pattern elements are similar to the model species, mimicry cannot be achieved successfully. In the Batesian mimicry butterflies, it is also known that not only wing patterns and shapes but also the flying behavior should resemble the model species (Kitamura and Imafuku 2005; Le Roy *et al*. 2019). In addition, some *Papilio* species show polymorphic Batesian mimicry, but few intermediate offspring between mimetic and non-mimetic types was observed (Clarke and Sheppard 1960; 1971; 1972; Clarke *et al*. 1968). These facts indicate that the multiple sets of traits for the mimicry are inherited in tightly linked manners, which have led to the “supergene” hypothesis (Clarke and Sheppard 1960). It was originally considered that supergene is composed of multiple flanking genes which are linked in the same chromosomal loci and inherited tightly together (Fisher 1930; Darlington and Mather 1949; Ford 1965; Hamilton 1964; Dobzhansky 1970). On the other hand, it has also been hypothesized that a single gene or a single regulatory element may regulate complex phenotypes such as mimicry by controlling multiple downstream genes (Nihout 1994; West-Eberhard 2003). Although many studies have reported that the supergene loci may be involved in the formation of complex phenotypes, no attempt has been made to reveal the functions of multiple genes within the supergene locus.

In a swallowtail *Papilio polytes*, only females have the mimetic and non-mimetic phenotypes, and males are monomorphic and non-mimetic (Fig. 1A). The mimetic female of *P. polytes* has red spots on the outer edge of hindwings and pale-yellow spots in the center of hindwings, which mimics the unpalatable model butterfly, *Pachliopta aristolochiae* (Euw *et al*. 1968; Clarke and Sheppard 1972; Uesugi 1996). The pale-yellow spots of the mimetic and non-mimetic forms differ not only in shape and arrangement, but also in the pigment composition (Nishikawa *et al*. 2013; Yoda *et al*. 2021). Males and non-mimetic females fluoresce under the UV irradiation, whereas mimetic females and model species, *Pachliopta aristolochiae*, do not fluoresce under the UV irradiation (Nishikawa *et al*. 2013; Yoda *et al*. 2021). It is also known that the mimetic female also resembles *Pachliopta aristolochiae* in the behavior of flight path (Kitamura and Imafuku 2005). Previous studies have shown that mimicry is regulated by the *H* locus and that the mimetic female (*H*) is dominant to the non-mimetic female (*h*) according to the Mendelian inheritance (Clarke and Sheppard 1972). Recently, whole genome sequences and genome-wide association studies have shown that about 130 kb of chromosome 25 which includes *doublesex* (*dsx*) is responsible for the *H* locus (Fig. 1B) (Kunte *et al*. 2014; Nishikawa et al. 2015). The direction of this region differs between the *H* allele and the *h* allele due to the inversions at both ends, suggesting that the *H* allele evolved from the *h* allele, given the conserved structure for the *h* allele type among lepidopteran insects (Kunte *et al*. 2014; Nishikawa et al. 2015). It is thought that recombination between the two alleles is suppressed by the inversion, and the accumulation of mutations and indels over the years has resulted in a highly diversified region (HDR) with low sequence homology between *H* and *h* (Nishikawa et al. 2015).

**Fig. 1.**
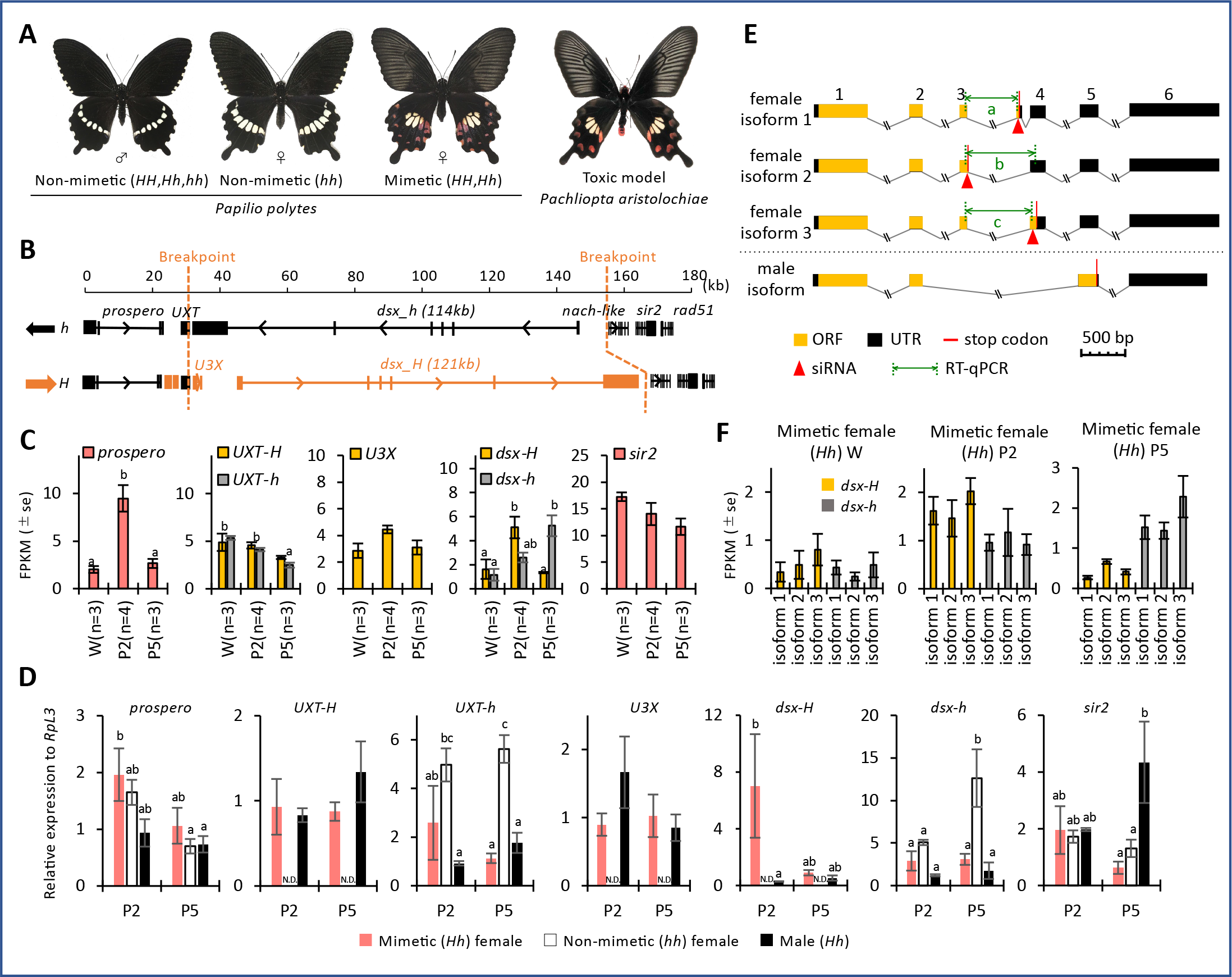
Wing patterns, structure of mimicry highly diversified region (HDR) and expression of genes within and flanking the HDR in *Papilio polytes*. (**A**) Wing patterns of adult male and non-mimetic and mimetic females of *P*. *polytes*, and toxic model, *Pachliopta aristolochiae*. Mimicry is regulated by *H* locus and mimetic allele (*H*) is dominant over the non-mimetic allele (*h*). (**B**) Detailed structure of mimicry HDR in *P*. *polytes* (Nishikawa *et al*. 2015,Iijima *et al*. 2018). The direction of the HDR is reversed between *h* and *H* (i.e., inversion). Putative breakpoints of the HDRs are indicated by orange dotted lines. The breakpoint on the left side is located inside the 5’-untranslated region (UTR) of the *ubiquitously expressed transcript* (*UXT*) gene, and that on the right side is located just on the outer side of *doublesex* (*dsx*). (**C**) Expression levels of genes within and flanking the HDR in hindwings of mimetic (*Hh*) females at the wandering stage (W) of the late last instar larvae, 2 day after pupation (P2) and 5 day after pupation (P5). FPKM values by RNA sequencing are shown with the error bars of standard error. Different letters indicate significant differences (Tukey post hoc test, *P*<0.05). (**D**) Relative expression levels of genes within and flanking the HDR in hindwings of mimetic (*Hh*) and non-mimetic (*hh*) females and males (*Hh*) at P2 and P5 estimated by RT-qPCR. *RpL3* was used as the internal control. Red, white, and black bars indicate mimetic and non-mimetic females and males, respectively. Error bars show standard error. Different letters indicate significant differences (Tukey post hoc test, *P*<0.05). (**E**) Schematic of *dsx* isoform in *P*. *polytes*. Female isoforms are divided into three types by start to stop codon sequences. There is one male isoform type. The region shown in black is the UTR, and the region shown in yellow is the open reading frame (ORF). Red bars indicate the position of the stop codon. The numbers above indicate the exon numbers (exon 1 to exon 6). Red arrowheads indicate the target of each isoform-specific siRNA, and green double arrows indicate the position of amplification by qPCR for quantification of each isofrom-specific expression. The sequences of siRNAs and primers are shown in Tables. S3 and S4. (**F**) Gene expression levels of each *dsx* isoforms in mimetic (*Hh*) females. FPKM values were calculated by RNA-seq at the wandering stage (W) of the late last instar larvae, at 2 day after pupation (P2) and at 5 day after pupation (P5). Orange bars indicate the expression levels of *dsx* isoforms from mimetic (*H*) allele and gray bars indicate from non-mimetic (*h*) allele. There was no statistically significant difference among isoforms.

Nishikawa *et al*. (2015) found in *P*. *polytes*, that knockdown of mimetic (*H*) type *dsx* (*dsx-H*) in the hindwings of mimetic females switched to a wing pattern similar to that of the non-mimetic females using the electroporation mediated RNAi method (Ando and Fujiwara 2013; Fujiwara and Nishikawa 2016). In addition, it is shown that *dsx-H* switches the pale-yellow colors from the UV fluorescent type (non-mimetic) to the UV reflecting type (mimetic), by repressing the papiliochrome II synthesis genes and nanostructural changes in wing scales (Yoda *et al*. 2021). Knockdown of non-mimetic (*h*) type *dsx* (*dsx-h*) did not cause such a switch, suggesting that *dsx-H* is essential for the formation of mimetic patterns, but the functional role of *dsx-h* is unknown (Nishikawa et al. 2015). The mimicry HDR contains not only *dsx* but also the 5’-untranslated region (UTR) portion of the *Ubiquitously Expressed Transcript* (*UXT*), a transcriptional regulator, and the long non-coding RNA *Untranslated 3 Exons* (*U3X*), present only in the HDR of the *H* allele (HDR-*H*) (Fig. 1B), but functions of *UXT* and *U3X* are still unclear (Nishikawa et al. 2015). In *P*. *memnon*, which is closely related to *P*. *polytes* and exhibits female-limited Batesian mimicry, the locus responsible for mimicry (*A*) is a *dsx*-containing region of chromosome 25 and consists of two types of HDRs with low homology between *A*-allele and *a*-allele (Komata et al. 2016; Iijima et al. 2018; Palmer and Kronforst 2020; Komata *et al*. 2022a). The mimetic-type HDR (HDR-*A*) of *P*. *memnon* also contains the 5’-UTR portion of *UXT* in addition to *dsx* (Iijima et al. 2018). Although an inversion is present in *P*. *polytes* and absent in *P*. *memnon*, the left-side breakpoint/boundary sites of the mimicry HDR is commonly located (Iijima et al. 2018). Furthermore, also in the closely related species, *P*. *rumanzovia*, which possesses the female-limited polymorphism, the left-side boundary of the mimicry HDR is thought to be located at the same position, i.e., in the 5’UTR of *UXT* (Palmer and Kronforst 2020; Komata et al. 2022b). These suggest that the left-side breakpoint/boundary sites of the mimicry HDR, i.e., the 5’ UTR of *UXT* and its surrounding regions, may have an important role in the regulation of the polymorphism (Komata et al. 2022a). Furthermore, in *P*. *polytes*, *UXT* and *U3X* are expressed in the hindwing, suggesting that these genes in the HDR may also be involved in the formation of mimetic patterns (Nishikawa *et al*. 2015).

Many supergenes show intraspecific polymorphism due to inversions, but in some cases, such as *P*. *memnon*, there is no inversion, but two types of HDR structures for mimetic and non-mimetic alleles are maintained (Iijima *et al*. 2018). However, it has not been clear whether the functional unit of the supergene that regulates complex adaptive traits is limited in the area within the inversion or the region of low homology (i.e., HDR), or whether it extends to neighboring regions. Both in *P*. *polytes and P. memnon*, the external gene *prospero*, which is adjacent to the internal gene *UXT* in the HDR, has read-through transcripts only in mimetic females (Nishikawa *et al*. 2015; Iijima *et al*. 2018). This suggests that some *cis*-regulatory element in the mimetic HDR may control the gene expression even in the external region, and that such a gene may be involved in the formation of the mimetic pattern.

In this study, we would like to elucidate the involvement of multiple genes other than *dsx-H* in the female-limited Batesian mimicry in *P*. *polytes* and the range of functional units in the supergene by examining the function of genes within and flanking the mimicry HDR. First, to search the allele- or phenotype- specific expression, the expression patterns of genes within and flanking the mimicry HDR were analyzed by RNA sequencing (RNA-seq) and reverse transcription quantitative PCR (RT-qPCR). Second, we explored the more detailed function of *dsx*: *dsx-H* is thought to switch mimetic and non-mimetic phenotypes, but the functional roles of *dsx-h* and three isoforms in *dsx* have been unclear. Third, to know the function of *UXT* and *U3X* other than *dsx* within the HDR-*H*, as well as *prospero* and *sir2* in close proximity to the outside of the HDR, we performed RNAi by *in vivo* electroporation (Ando and Fujiwara 2013; Fujiwara and Nishikawa 2016) and Crispr/Cas9 knockout for *UXT*. In addition, RNA-seq was performed on the *dsx-H*, *UXT*, and *U3X* gene knockdown wings and the control wings to elucidate the regulatory relationship and downstream genes of the three genes.

## Materials and Methods

### Butterfly rearing

We purchased wild-caught *P*. *polytes* from Mr. Y. Irino (Okinawa, Japan) and Mr. I. Aoki (Okinawa, Japan), and obtained eggs and used for the experiment. The larvae were fed on the leaves of *Citrus hassaku* (Rutaceae) or on an artificial diet, and were kept at 25 °C under long-day conditions (light:dark = 16:8 h). Adults were fed on a sports drink (Calpis, Asahi. Japan).

### Analysis of expression levels of genes in and flanking the HDR

In this study, we used the entire hindwing of *P*. *polytes* to analyze the expression levels of internal (*U3X*, *UXT*, *dsx*) and external flanking genes (*nach-like*, *sir2*, *prospero*) in the HDR by RNA-seq and RT-qPCR. In addition to the published RNA-seq read data of *P*. *polytes* (BioProject ID: PRJDB2955) (Nishikawa *et al*. 2015), newly sampled RNA was used for the analysis. The sample list used in the experiment is shown in Table S1. The RNA-seq data used in this study include both data obtained in previous studies and newly obtained data in this study, but since the data were obtained in the same laboratory using the same method and the trends of expression levels are consistent with those by RT-qPCR results described below, correction for the Batch effect was not performed. The newly added RNA-seq reads were obtained by the following procedure. The entire hindwing was sampled for RNA extraction on pupal day 2 (P2) and pupal day 5 (P5), and RNA extraction was performed using TRI reagent (Sigma) in the same manner as Nishikawa et al. (2015) and Iijima et al. (2019). The extracted and DNase I (TaKaRa, Japan) treated RNA was sent to Macrogen Japan Corporation for library preparation by TruSeq stranded mRNA (paired-end, 101 bp) and sequenced by Illumina platform. The obtained RNA-seq reads were quality-checked by FastQC (version 0.11.9) (Andrews 2010), mapped by Bowtie 2 (version 2.4.4) (Langmead and Salzberg 2012), and the number of reads was counted using SAMtools-(version 1.14) (Li *et al*. 2009). Based on the number of reads, FPKM (Fragments Per Kilobase of transcript per Million mapped reads) was calculated (FPKM=number of mapped reads/gene length(bp)/total number of reads×10^9^). For mapping, full-length mRNA sequences including UTRs were used for *prospero*, *UXT-H*, *UXT-h*, *U3X*, *sir2*, and *nach-like*, and ORF region sequences were used for *dsx-H* and *dsx-h*. *dsx-H* and *dsx-h* were mapped to three female isoforms for female individuals and one male isoform for male individuals. Sequence information for each gene was obtained from Nishikawa et al. (2015) and Iijima et al. (2018).

The expression levels of *U3X*, *UXT-H*, *UXT-h*, *dsx-H*, *dsx-h*, *sir2*, and *prospero* were also analyzed by RT-qPCR. RNA obtained by the above method was subjected to cDNA synthesis using Verso cDNA synthesis kit (Thermo Fisher Scientific). The qPCR was performed using Power SYBR® Green Master Mix (Thermo Fisher Scientific) by QuantStudio 3 (ABI). The detailed method was followed by Iijima et al. (2019). A total of 18 whole hindwing samples from 18 individuals were used, including three each of mimetic females (*Hh*), non-mimetic females (*hh*), and males (*Hh*) of P2 and P5. *RpL3* was used as an internal standard and the primers used are shown in Table S3.

### Knockout of *UXT* by Crispr/Cas9

A single guide RNA (sgRNA) was used to generate deletions and frameshifts within the prefoldin domain of *UXT* (Figure S7). A sgRNA was designed using CRISPRdirect (https://crispr.dbcls.jp), and the specificity of the sequence of sgRNA was assessed using BLAST to ensure that there were no multiple binding sites. The target sequence is shown in Figure S7. The sgRNA template was generated by PCR amplification with forward primers encoding the T7 polymerase binding site and the sgRNA target site (Pp_UXT_F1modi, GAAATTAATACGACTCACTATAGGCCGACCAGAAGCTTCATCGTTTAAGAGCTATGCTGGAAACAGCATAGC), and reverse primers encoding the remainder of the sgRNA sequence (sgRNA_Rmodi, AAAAGCACCGACTCGGTGCCACTTTTTCAAGTTGATAACGGACTAGCCTTATTTAAACTTGCTATGCTGTTTCCAGCATA), using Phusion DNA polymerase (M0530, New England Biolabs, Ipswich, MA, USA) (Zhang and Reed 2017). *In vitro* transcription was performed using the Megascript T7 Kit (Thermo Fisher Scientific) and sgRNA was purified with the MEGAclear Transcription Clean-Up Kit (AM1908, Thermo Fisher Scientific). To collect eggs for injection, host plants were provided to female butterflies and allowed to lay eggs for 1 hour. The obtained eggs were aligned on a glass slide and fixed with an instant glue Aron Alpha (Toagosei Company, Japan). The fixed eggs were disinfected with formalin for 3 min, the tip of the glass capillary was cut with a razor at an angle of 30–40°, perforated with a tungsten needle, and the capillary was injected with an injection mixture containing sgRNA (500 ng/ul) and Cas9 protein (CP-01, PNA Bio; 500 ng/ul) (injection pressure Pi 100 Pa, steady pressure Pc 40–80 Pa). Finally, the holes were sealed with Arone Alpha, placed in a Petri dish, and stored in a plastic case along with a well-moistened comfort towel. The hatched larvae were reared in the same manner as described above. The emerged adults were observed for phenotype, and parts of the head, abdomen and wings were taken for genotyping. DNA was extracted using a phenol-chloroform protocol and PCR amplified across the target sites (primers, Pp_UXT_cr_F1, ttcgtgttcaggatcaacag; Pp_UXT_cr_R1, tatttgttaactgcccgatg). PCR products were used to perform TA cloning, Sanger sequencing, and genotyping.

### Functional analysis by RNAi using *in vivo* electroporation

siDirect (http://sidirect2.rnai.jp/) was used to design the siRNAs. The target sequences were blasted against the predicted gene sequence (BioProject: PRJDB2954) and the genome sequence (BioProject: PRJDB2954) in *P*. *polytes* to confirm that the sequences were highly specific, especially for the target genes. The designed siRNA was synthesized by FASMAC Co., Ltd. (Kanagawa, Japan). The RNA powder received was dissolved in Nuclease-Free Water (Thermo Fisher, Ambion), adjusted to 500 μM, and stored at −20°C. The sequence information of the siRNA used is listed in Table S4. As a negative control, universal negative control siRNA (Nippongene), designed to target sequences that were found not to show homology with all eukaryotic genes, was adjusted to 250 μM. A glass capillary (Narishige, GD-1 Model, Glass Capillary with Filament) was processed into a needle shape by heating it at HEATER LEVEL 66.6 using a puller (Narishige, PP-830 Model). The capillary was filled with siRNA. siRNA was adjusted to 250 μM when only one type of siRNA was used for one target gene (*dsx-H*, *dsx-H* female isoform 1–3, *dsx-h*&*H*, *UXT*, *U3X*, *rn*), and 500 μM siRNA solution was mixed in equal amounts when two types of siRNA were mixed for one target gene (*prospero*, *sir2*). For all genes except *U3X*, siRNAs were designed in the ORF. Since *UXT* differs from *UXT-H* and *UXT-h* sequences only in the 5’UTR, the knockdown in this experiment was targeted to the common ORF region of *UXT-H* and *UXT-h*. The capillary was filled with siRNA and 4 μl of siRNA was injected into the left hindwing under a stereomicroscope using a microinjector (FemtoJet, eppendorf). Then, siRNA was introduced into only the positive pole side of the electrode by applying voltage (5 square pulses of 7.5 V, 280 ms width) using an electroporator (Cellproduce, electrical pulse generator CureGine). A PBS gel (20×PBS: NaCl 80g, Na2HPO4 11g, KCl 2g, K2HPO4 2g, DDW 500ml; 1% agarose) was placed on the dorsal side of the hindwing and a drop of PBS was placed on the ventral side of the hindwing. The detailed method follows that described in the previous paper (Ando and Fujiwara 2013). The pictures of all the individuals who performed the function analysis are described collectively as Supplementary figures.

### Regulatory relationship of *dsx-H*, *UXT*, and *U3X* expression by RNAi and downstream gene screening

After sampling the hindwings of individuals with *dsx-H*, *UXT*, and *U3X* knockdown by RNAi in the P2 stage with the siRNA-injected side as knockdown and the non-injected side as control, total RNA was extracted and DNase I treated RNA was sent to Macrogen Japan Corporation. Libraries were prepared using TruSeq stranded mRNA (paired-end, 101 bp) and sequenced using the Illumina platform. Sample and read information are shown in Table S2. dsx-H_Control_2 and dsx-H_knockdown_2 are the read data used in a previous study (Iijima *et al*. 2019). A portion of the total RNA used for RNA-seq was used for cDNA synthesis, and RT-qPCR was used to pre-confirm the reduced expression of the knockdown side. For RNA-seq analysis, we first performed quality check using FastQC (Version 0.11.9) (Andrews 2010), and the reads were mapped to the transcript sequences of *P*. *polytes* to calculate the expression levels. The transcript sequence was obtained from NCBI, GCF_000836215.1_Ppol_1.0_rna.fna (BioProjects: PRJNA291535, PRJDB2954). Because the transcript sequence information of the genes around *H* locus described in GCF_000836215.1_Ppol_1.0_rna.fna was incomplete (*H* and *h* derived transcripts of *dsx* and *UXT* were confused, and especially for *dsx*, incomplete transcripts containing *dsx* fragments and male isoforms are included.), read mapping to the genes around *H* locus (*prospero*, *UXT*, *U3X*, *dsx-H*, *dsx-h*, *sir2*, *rad51*) was performed separately: the full-length mRNA sequences including UTRs were used for *prospero*, *UXT*, *U3X*, *sir2*, and *rad51*, and the ORF region sequences of female isoforms for *dsx-H* and *dsx-h* were used. Mapping and calculation of FPKM value were performed as described above.

In addition, R software was used to extract genes with variable expression by statistical analysis of read data, and comparison between two groups with correspondence using Wald-test of DESeq2 (version 3.14) (Love *et al*. 2014) was performed. The transcription factors and signaling factors were extracted using the GO terms of the top hit amino acid sequences by Blastx against the Uniprot protein database. “DNA-binding Transcription factor activity” [GO:0003700], “DNA-binding transcription factor activity, RNA polymerase II-specific” [GO:0000981] as transcription factors and “signaling receptor binding” [GO:0005102], “signaling receptor activity” [GO:0038023] as signaling factor.

Finally, using RNA-seq data from three mimetic and three non-mimetic females of P5 each (Table S1), genes whose expression were elevated in mimetic and non-mimetic females were extracted using the same method described above.

### Statistical analysis

Statistical analysis of the data was performed with R software (R Core Team 2020). In the analysis of gene expression levels (Figs. 1, C, D and F), we explored the effects of stage and/or genotype/sex using a generalized linear model (GLM) with a normal distribution. Tukey’s post hoc tests were used to detect differences between groups using the “glht” function in the R package multcomp (Hothorn *et al*. 2008); *P*<0.05 was considered statistically significant. For the analysis to examine the effects of gene knockdown (Figs. 3, D–F, 6A and S13), paired *t*-test was used; *P*<0.05 was considered statistically significant.

## Results

### Comparison of the expression levels of genes inside and flanking the mimicry HDR-*H*

It is reported that wing coloration occurs after day 9 of pupation (P9), and that the expression of *dsx-H* peaked on days 1–3 of pupation (P1–P3) (Nishikawa *et al*. 2013; 2015). Therefore, although more detailed studies are needed to determine when the important period for the function of *dsx-H* is, we focused on gene expression in the first half of the pupal stage to investigate the mimetic wing pattern formation. In order to investigate whether each gene in and flanking the HDR-*H* is involved in these processes, we examined the expression levels of the genes, *prospero*, *UXT-H* (*UXT* from *H*-allele), *UXT-h* (*UXT* from *h*-allele), *U3X*, *dsx-H*, *dsx-h*, *nach-like*, and *sir2* (Table 1 shows the summary of gene functions.), in the hindwing imaginal discs at the wandering stage (W) of the last instar larvae, P2 and P5, by RNA-seq (Table S1 shows the list of samples used). The results showed that *nach-like* was not expressed at all in any developmental stage as reported in *P*. *memnon* (Iijima *et al*. 2018). Other genes were expressed in mimetic females, non-mimetic females and males (Figs. 1C and S1).

**Table 1.**
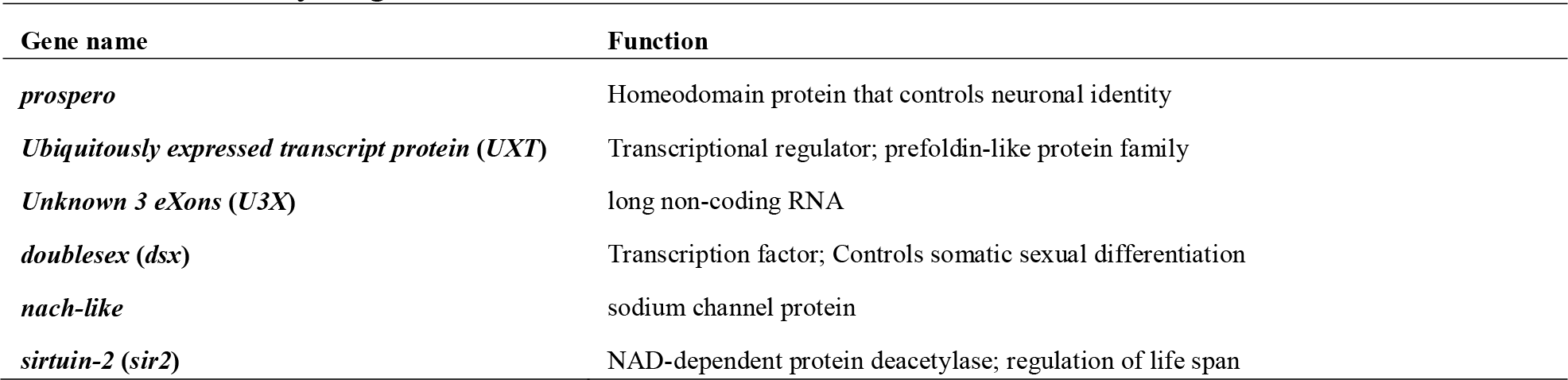
Summary of gene function in/around HDR.

In the RNA-seq data of mimetic females, *dsx-H* and *dsx-h* showed contrasting expression patterns. *dsx-H* showed a peak expression in the P2 stage, while *dsx-h* was highly expressed in the P5 stage (Fig. 1C). The expression of *U3X*, which is only present in the *H* locus, tended to show the constant expression and relatively high in P2, while the data was not statistically significant (Fig. 1C). There was no significant difference in the expression pattern of *UXT* between *UXT-H* and *UXT-h*, and the expression level of *UXT* was higher in W and P2 and significantly lower in P5 (Fig. 1C). The expression pattern of *prospero* was significantly larger in P2 as in *dsx-H*, and that of *sir2* was largest in W as in *UXT*, but not statistically significant (Fig. 1C), In the RNA-seq experiment, we used three or more samples at each stage (W, P2, P5) for mimetic females, but insufficient numbers of samples for some non-mimetic females and males (Fig. S1), and thus we further performed RT-qPCR using P2 and P5 samples for mimetic females (*Hh*), non-mimetic females (*hh*), and males (*Hh*) (Fig. 1D).

RT-qPCR showed that the *dsx-H* expression was significantly high in P2 of mimetic females but low in P5 and males in every stage (Fig. 1D). The *dsx-h* expression was significantly high in P5 of non-mimetic females compared to other stages, mimetic females and males (Fig. 1D). It is noteworthy that the expression of *dsx* was low in males at all stages (Fig. 1D). The high expression of *dsx-H* in P2 of mimetic females is consistent with RNA-seq results (Fig. 1C) and previous studies, which may be related to the mimetic color pattern formation (Nishikawa *et al*. 2015; Deshmukh *et al*. 2020). Expression of *dsx-h* in mimetic females is also consistent with RNA-seq results; RNA-seq results show significantly greater expression in P5 than in W, but comparing P2 and P5, P5 tends to be larger but not significantly different. RT-qPCR showed no significant difference in the expression of *dsx-h* in mimetic females because we measured the expression only in P2 and P5, then more detailed expression analysis is needed to elucidate more detailed expression patterns of *dsx-h*. *U3X* was not detected in non-mimetic females (*hh*) because it is present only in the *H* locus, and was expressed in mimetic females and males at P2 and P5 stages (Fig. 1D). For *UXT-H*, there was no significant difference in expression levels among mimetic females and males in any stages (Fig. 1D). The expression of *UXT-h* was significantly greater in non-mimetic females (*hh*), probably because they are *h* homozygous, but it was particularly high in non-mimetic females in P5, about five times higher than the expression of *UXT-h* in mimetic females (Fig. 1D). The expression of *sir2* and *prospero* in mimetic females was similar to that of RNA-seq results, and their expression was also observed in non-mimetic females and males (Fig. 1D). The expression of *sir2* was significantly higher in males in P5 than in mimetic or non-mimetic females (Fig. 1D).

To summarize the results of the expression analyses by RNA-seq and RT-qPCR, *dsx-H* and *dsx-h* appear to be regulated separately, as *dsx-H* and *dsx-h* showed contrasting expression patterns. The trend of high expression at P2 as well as *dsx-H* was observed in *prospero* and *U3X*, and the trend of gradually decreasing expression at W, P2, and P5 was observed in *UXT-H*, *UXT-h*, and *sir2*. *prospero* may be regulated by genes/genetic elements in the mimicry HDR, as its expression pattern is the same as that of *U3X* and *dsx-H*, which are specific to the *H* allele. We did not find any genes with similar expression patterns to *dsx-h*. More detailed spatial-temporal expression patterns will need to be examined with related genes in the future, as it is possible that the gene expression is elevated only at specific spots or stages and contributes to the formation of color patterns and spots in butterfly wings.

### Expression and functional roles of *dsx-h* and three isoforms of *dsx*

It is important to clarify the functional roles of *dsx-H* and *dsx-h* in the evolution of mimicry supergene. Although the involvement of *dsx-H* in the formation of mimetic traits has been shown, the function of *dsx-h* has been unclear (Nishikawa *et al*. 2015; Iijima *et al*. 2019; Yoda *et al*. 2021). The hindwing patterns of non-mimetic females and males have small differences in the bright field (only the non-mimetic females have blue and red spots blow the innermost pale-yellow spot), but are clearly distinguishable when observed under the UV irradiation (see Nontreated in Fig. 2, B and C). In non-mimetic females, the innermost and second innermost pale-yellow spots do not fluoresce, whereas in males the innermost one fluoresces slightly and the second one fluoresces completely. We injected siRNA of the target gene (i.e., *dsx* in this case) into the hindwing immediately after pupation (Day 0 of pupation: P0) and performed electroporation to cover most of the hindwing, which induces RNAi only in the target area (Ando and Fujiwara 2013; Fujiwara and Nishikawa 2016). Although this method has already been established in previous studies (Nishikawa et al. 2015; Iijima et al. 2019) and can be used for experiments, we introduced Universal Negative Control siRNA (Nippongene) as a negative control and confirmed that there was no phenotypic change (Figs. 2A and S2). When *dsx* was knocked down in non-mimetic females (*hh*), the second pale-yellow spot fluoresced, showing a similar pattern to males (Figs. 2B and S3). The difference between UV fluorescence and reflection is thought to be caused by the difference in whether the pigment papiliochrome II is synthesized or not (Yoda et al. 2021). Therefore, the confirmation of the UV fluorescence at the second pale-yellow spot by knockdown of *dsx-h* in non-mimetic females is thought to be due to the fact that papilochrome II is synthesized by knockdown of *dsx-h*. In males, however, there was no clear change after knockdown of *dsx* (Figs. 2C and S3), indicating that although the male isoform of *dsx* has no function in male wing color pattern, *dsx-h* maintains its original function of sexual differentiation in the hindwing as well as the mimetic pattern formation.

**Fig. 2.**
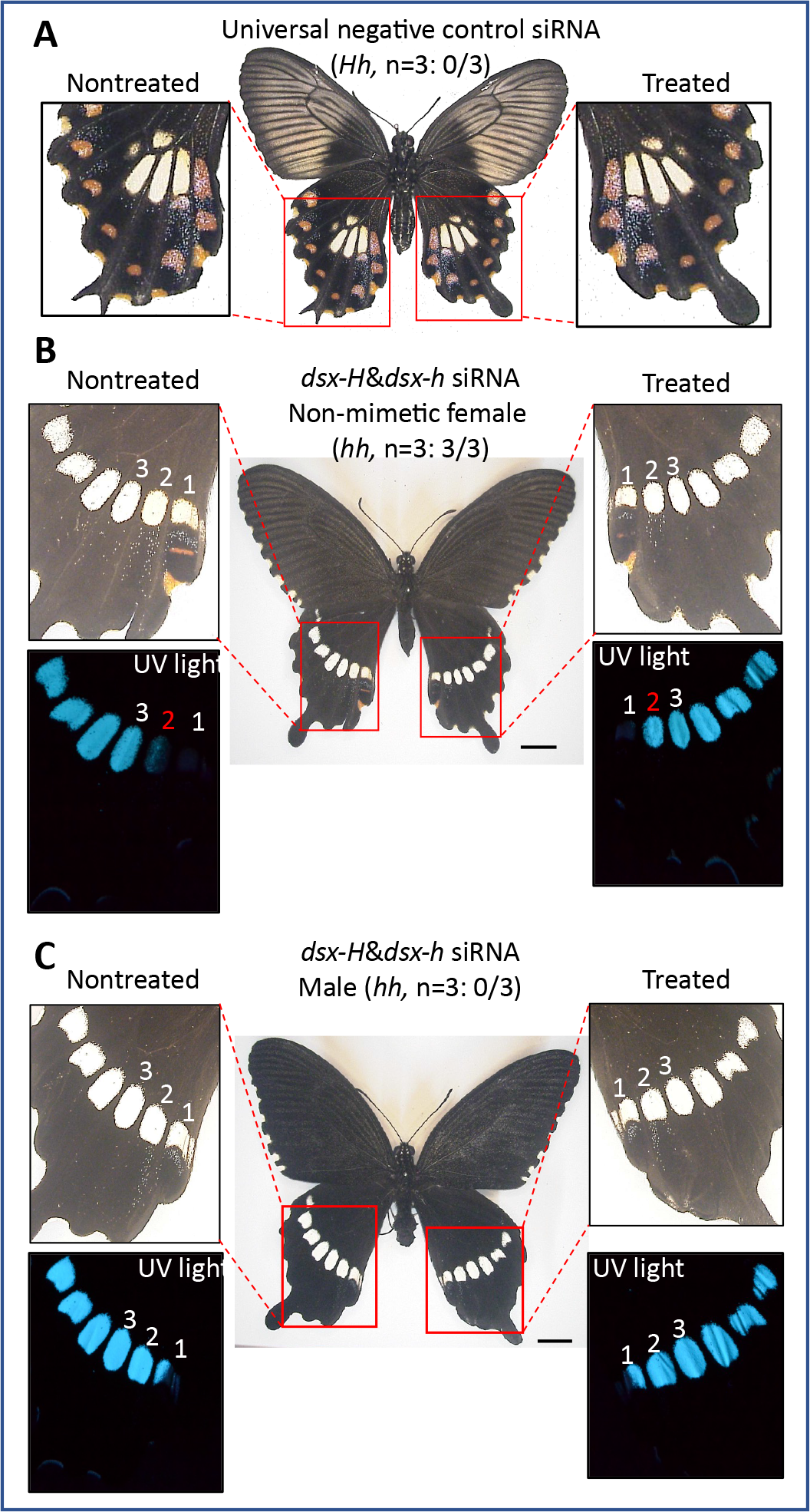
Functional analyses of *dsx* in non-mimetic females and males of *Papilio polytes*. **(A)** A negative control using Universal Negative Control siRNA (Nippongene) designed to target sequences that were found not to show homology with all eukaryotic genes. No phenotypic effects were detected. (**B, C**) Knockdown of *dsx-H*&*dsx-h* in the left hindwings of non-mimetic (*hh*) female (**B**) and male (**C**). The siRNA targeting sequence common to all alleles and isoforms of *dsx* was injected into the left pupal hindwing immediately after pupation and electroporated into the dorsal side. Pale-yellow spots are numbered starting from the inside. The numbered spots in red indicate those whose UV response was altered by knockdown. In non-mimetic females (**B**), knockdown of *dsx* changed the second spot to produce UV fluorescence like males (*hh*) **(C)**. Scale bars, 1cm. *dsx* genotypes and the number of samples showing phenotypic changes among the tested numbers are shown in parentheses Supplementary Figure S3 show other replicates.

**Fig. 3.**
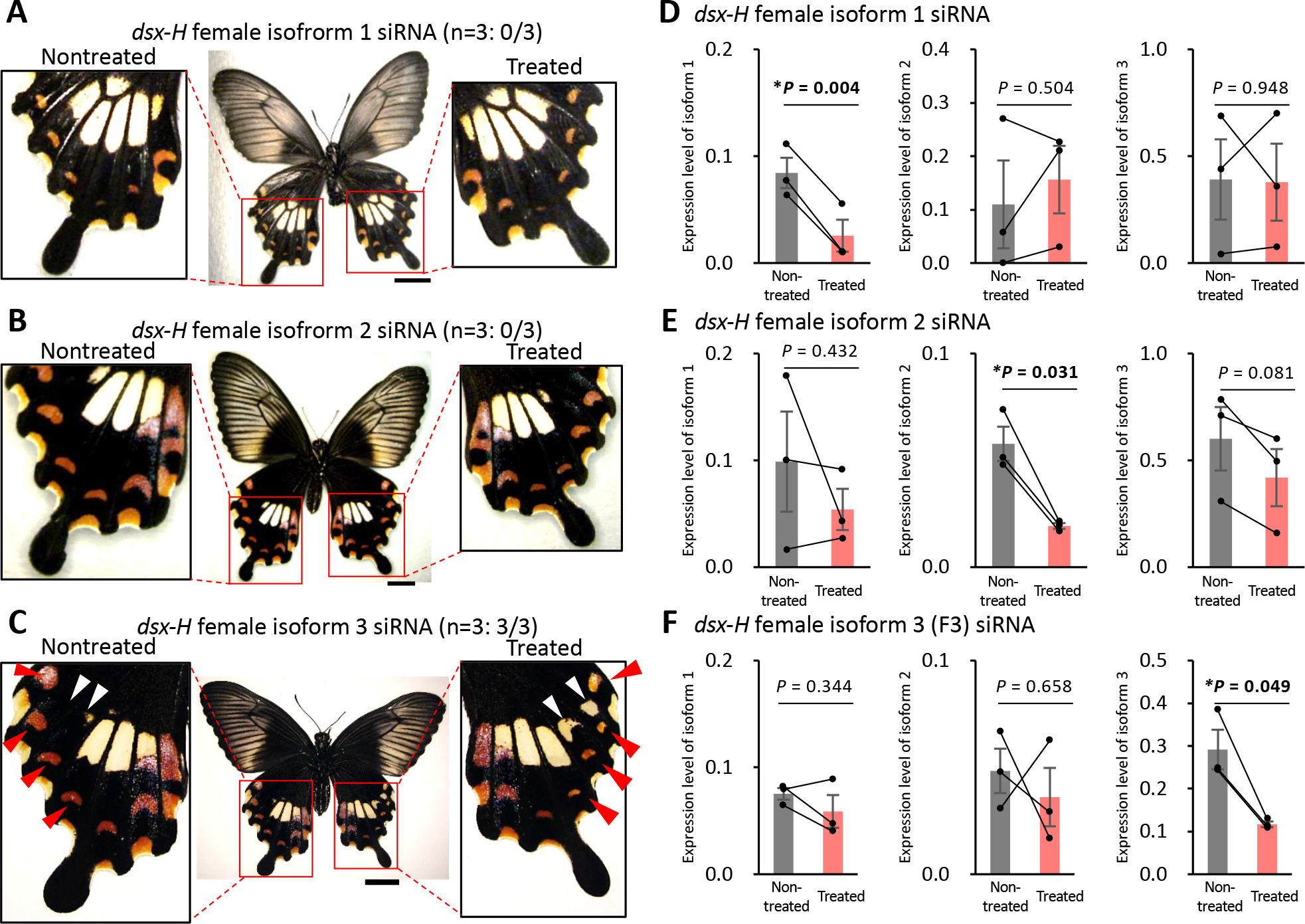
Knockdown experiments of *dsx-H* female isoform 1, 2 and 3 in the hindwings of mimetic (*Hh*) females. siRNA was injected into the left pupal hindwing immediately after pupation and electroporated into the ventral side. No phenotypic changes were observed by knockdowns of *dsx-H* female isoform 1 and 2 (**A, B**), but knockdown of *dsx-H* female isoform 3 changed the mimetic pattern to the similar of the non-mimetic pattern (**C**). Red and white arrowheads represent the changed red and pale-yellow regions, respectively. Scale bars, 1 cm. The number of samples showing phenotypic changes among the tested numbers is shown in parentheses. Supplementary Figure S5 show other replicates. (**D**–**F**) Gene expression levels of each isoform in the knockdown wings of mimetic (*Hh*) females at 2 days after pupation. When *dsx-H* female isoform 1 was knocked down, only *dsx-H* female isoform 1 was down-regulated (**D**), when *dsx-H* female isofom 2 was knocked down, only *dsx-H* female isoform 2 was down-regulated (**E**), and when *dsx-H* female isofom 3 was knocked down, only *dsx-H* female isoform 3 was down-regulated (**F**). Gray and red bars shown the expression in nontreated and treated hindwings, respectively. We estimated the gene expression levels by RT-qPCR using *RpL3* as the internal control. Error bars show standard error of three biological replicates. \**P* < 0.05 for paired t-test.

In addition, there are three female isoforms (F1, F2, F3) both in *dsx-H* and *dsx-h*, and one isoform in *dsx-H* and *dsx-h* in males (Fig. 1E) (Kunte *et al*. 2014;Nishikawa *et al*. 2015). The role of these isoforms is scarcely known in *P*. *polytes* or in other insects. To investigate the function of the three isoforms in females, we performed expression analysis by RNA-seq and knockdown experiment by RNAi. ORF region sequences were used for mapping to three female isoforms in each of *dsx-H* and *dsx-h* for female individuals and one male isoform in each of *dsx-H* and *dsx-h* for male individuals. There was no significant difference in the expression levels of F1, F2, and F3, but F3 showed relatively higher expression levels (Figs. 1F and S4). Next, we designed isoform specific siRNAs and primers (Fig. 1E and Tables. S3 and S4). RNAi by *in vivo* electroporation showed that only the F3-specific knockdown of *dsx-H* changed the mimetic pattern to the similar of the non-mimetic pattern (Figs. 3, A–C and S5): red spots became smaller, and the non-mimetic specific pale-yellow spots appeared (Figs. 3C and S5). In the RNAi experiment, we confirmed that only the target isoform was down-regulated by RT-qPCR (Fig. 3, D–F). Although the function of the three female isoforms of *dsx-h* was not studied, these results indicate that only isoform 3 of *dsx-H* has an important function in the formation of mimetic traits.

### Functional analysis of *UXT* and *U3X* in the HDR

*UXT* is a gene in the prefoldin-like protein family of transcriptional regulators whose 5’UTR is contained in the HDR. In the *UXT* knockdown by siRNA injection in the hindwing, the pale-yellow spots were reduced, and the shape of pale-yellow region was flattened like non-mimetic phenotype, and the red spots were reduced or disappeared (Figs. 4A and S6). But the UV response in the pale-yellow spots was not changed by the *UXT* knockdown (Fig. S6). Knockout of *UXT* by Crispr/Cas9 was also performed. guide RNA was designed to target the functional domain of *UXT*, the prefoldin domain (Fig. S7), and was injected into eggs immediately after egg laying together with Cas9 protein (CP-01, PNA Bio). A total of 294 eggs were injected and 21 adults were obtained (Mimetic female: 8; nonmimetic female: 5; male: 8, Fig. S7). Eight of the mimetic females were subjected to PCR, cloning, Sanger sequencing to confirm the introduction of mutations (Fig. S8), and phenotypic observation. The adult eclosionrate (7.1%) of the injected individuals was very low, suggesting that *UXT* may have an important function in survival. Genotyping using DNA extracted from the abdomen, head and wings of emerged individuals yielded five types of sequences in which mutations were introduced (Fig. S8). We observed individuals with the mosaic knockout, in which the pale-yellow spots were flattened as in the non-mimetic form, and the red spots were reduced or disappeared (Fig. 4B). In only one individual, the pale-yellow spots were changed, and the red spots were reduced in four individuals (Fig. S9). The UV response in the pale-yellow spots was not changed in any individuals (Fig. S9). These results indicate that *UXT* is involved in mimetic pattern formation in both pale-yellow and red spots. In pale-yellow spots, both *dsx-H* and *UXT* are involved in the mimetic pattern formation, but *dsx-H* acts to suppress the two pale-yellow spots characteristic of non-mimetic female in outer side of hindwings (Fig. 3C) and switches the pale-yellow colors from the UV fluorescent type (nonmimetic) to the UV reflecting type (mimetic) (Yoda et al. 2021), whereas *UXT* is thought to be involved in the overall shape of the pale-yellow spots (Fig. 4, A and C).

**Fig. 4.**
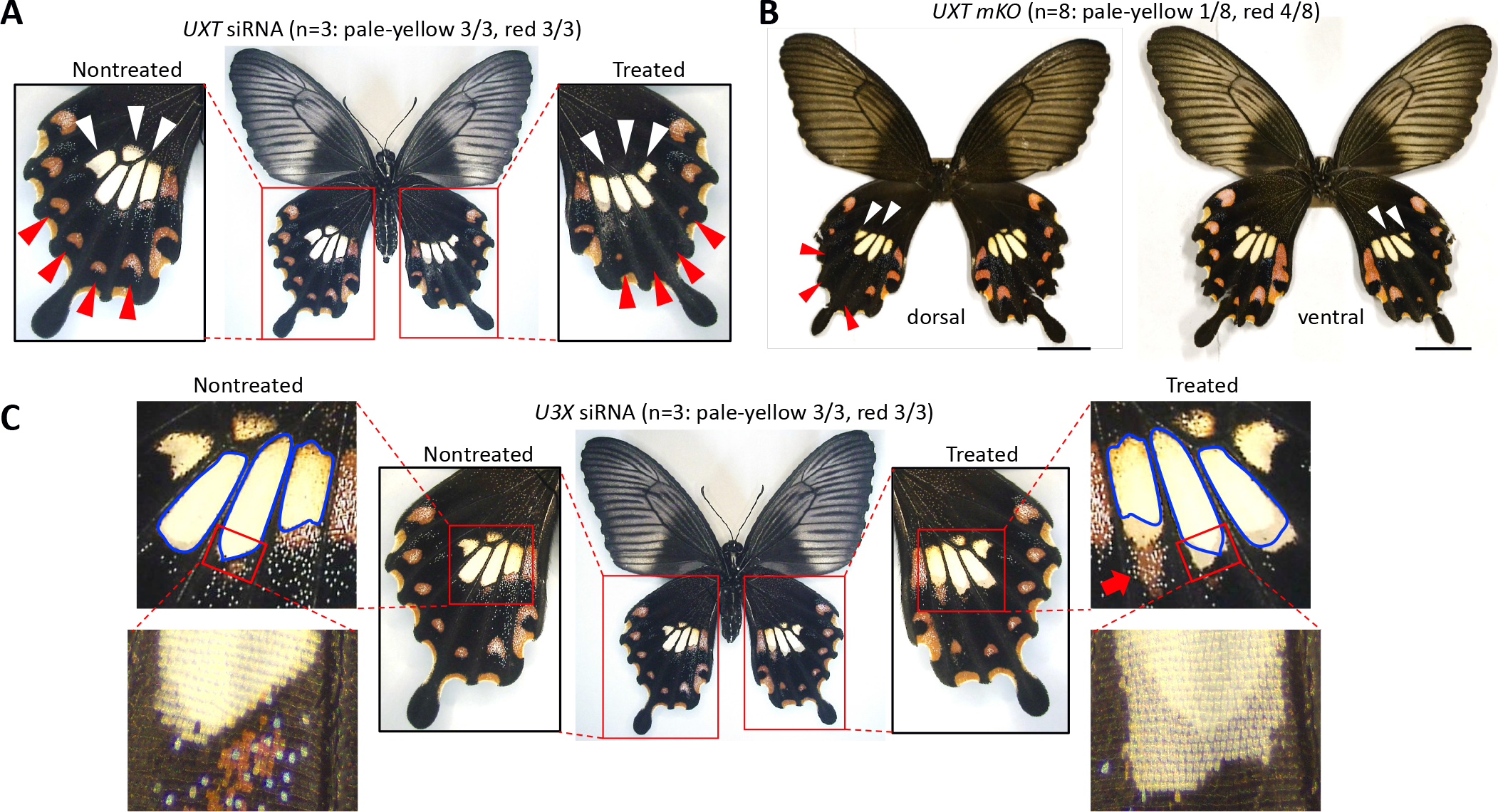
Functional analysis of *UXT* and *U3X* in *Papilio polytes*. (**A**) Knockdown of *UXT* in the hindwings of mimetic (*Hh*) females. siRNA was injected into the left pupal hindwing immediately after pupation and electroporated into the ventral side. Red and white arrowheads represent the changed red and pale-yellow regions, respectively. Supplementary Figures S6 show other replicates. (**B**) Mosaic knockout of *UXT* by Crispr/Cas9. Dorsal and ventral views of one representative of the eight individuals observed are shown. Red and white arrowheads represent the changed red and pale-yellow regions, respectively. In this individual, phenotypic changes were observed mainly on the left hindwing in dorsal view. Scale bars, 1 cm. Supplementary Figure S9 shows other replicates. (**C**) Knockdown of *U3X* in the hindwings of mimetic (*Hh*) females. The red arrow indicates the area where the red spot has expanded, and the area circled by blue line indicates the original area of pale-yellow spots. On the treated side, the area of pale-yellow spots has slightly extended. Further magnification of the enlarged area of pale-yellow spots shows that only on one side of the siRNA-intruduced hindwing, the original black scales have changed to pale-yellow scales. The number of samples showing phenotypic changes among the tested numbers is shown in parentheses. Supplementary Figures S10 show other replicates.

In the *U3X*, which is long non-coding RNA, knockdown, the pale-yellow spots were extended downward (Figs. 4C and S10), and the red spots below the innermost pale-yellow spots, which were only slightly visible in the control, were enlarged (indicated by red arrow in Fig. 4C). Phenotypic changes were observed by *U3X* knockdown, but not simply a change from mimetic to non-mimetic phenotype. Since *U3X* is a non-coding RNA, the phenotypic changes observed upon knockdown of *U3X* may be due to changes in the expression of other genes that are regulated by *U3X*. The expansion of red and pale-yellow spots upon knockdown of *U3X* suggests the existence of genes that play a role in suppressing the excessive appearance of these spots.

### Functional analysis of *prospero* and *sir2*, two proximal genes outside the HDR-*H*

Next, the functional roles of *prospero* and *sir2* which locate in close proximity to the HDR region but outside the inversion, were analyzed in mimetic female hindwings by *in vivo* electroporation mediated RNAi. In the *prospero* siRNA injected hindwing, there were no obvious changes, but the red spots characteristic of the mimetic form were subtly enlarged in three of the six individuals tested (Figs. 5A and S11). Although in the *prospero* knockdown the red spot was enlarged in half of the individuals tested, it may be difficult to definitively conclude the function of *prospero* because the red spots were reduced in some individuals (Fig. S11). However, it is probably true that *prospero* is involved in the formation of red spots. In the case of *sir2* RNAi, the pale-yellow spots were flattened like non-mimetic phenotype in four of the seven individuals tested, and the red spots were enlarged under the innermost pale-yellow spots in five individuals (Figs. 5B and S12). The decreases in *prospero* and *sir2* expressions by RNAi were confirmed by RT-qPCR, and although not statistically significant, there was a tendency for those expressions to decrease in the knockdown side (Fig. S13). These results suggest that multiple adjacent genes outside the HDR are involved in the formation of mimetic patterns.

**Fig. 5.**
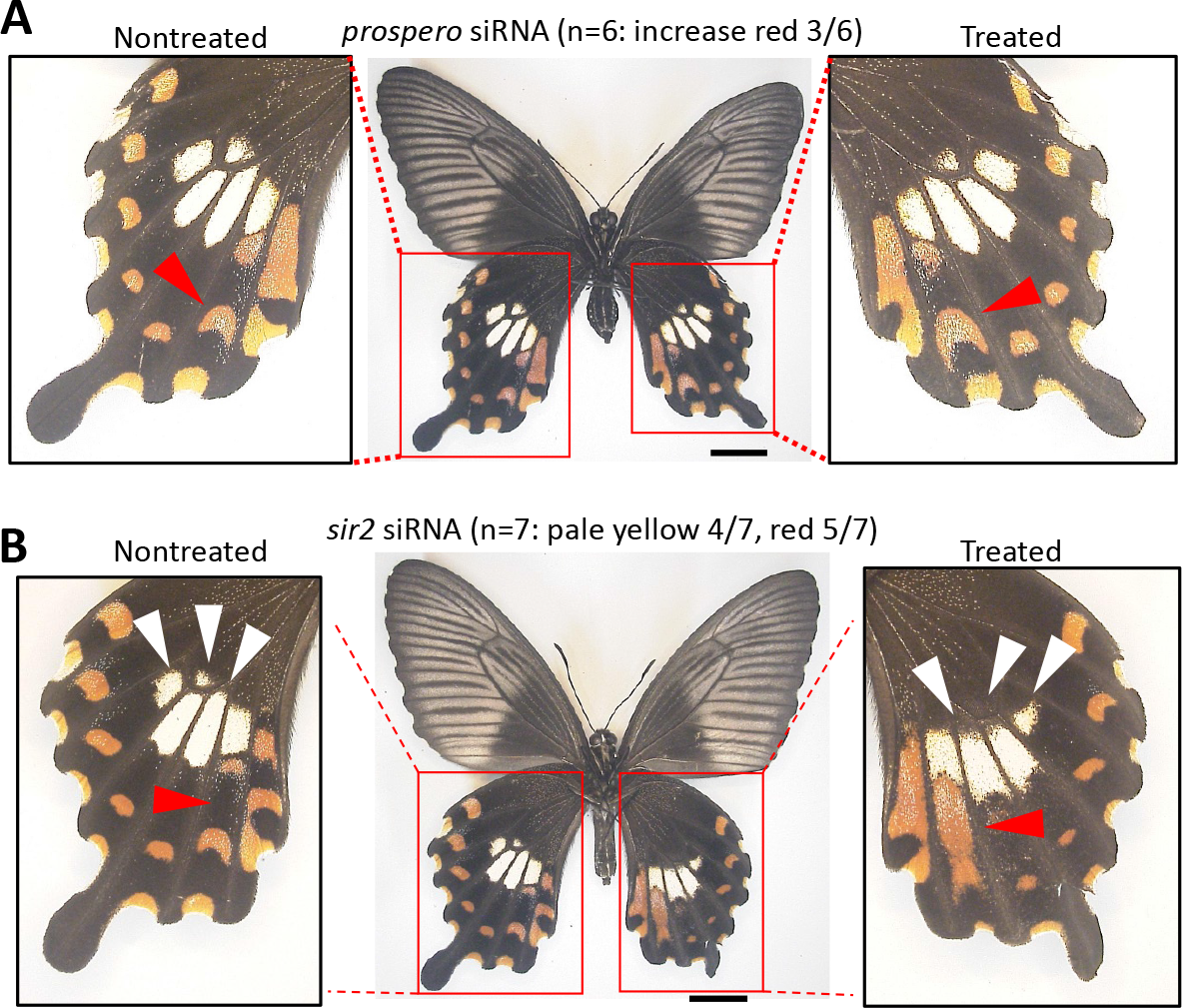
Knockdown of *prospero* **(A)** and *sir2* **(B)** in the hindwings of mimetic (*Hh*) females of *Papilio polytes*. siRNA was injected into the left pupal hindwing immediately after pupation and electroporated into the ventral side. The red spots characteristic of the mimetic form were subtly enlarged in *prospero* knockdown (**A**). In the *sir2* knockdown, the pale-yellow spots were flattened like non-mimetic phenotype, and the red spots were enlarged under the innermost pale-yellow spots (**B**). Red and white arrowheads represent the changed red and pale-yellow regions, respectively. Scale bars, 1cm. The number of samples showing phenotypic changes among the tested numbers is shown in parentheses. Supplementary Figs. S11, 12 show other replicates.

**Fig. 6.**
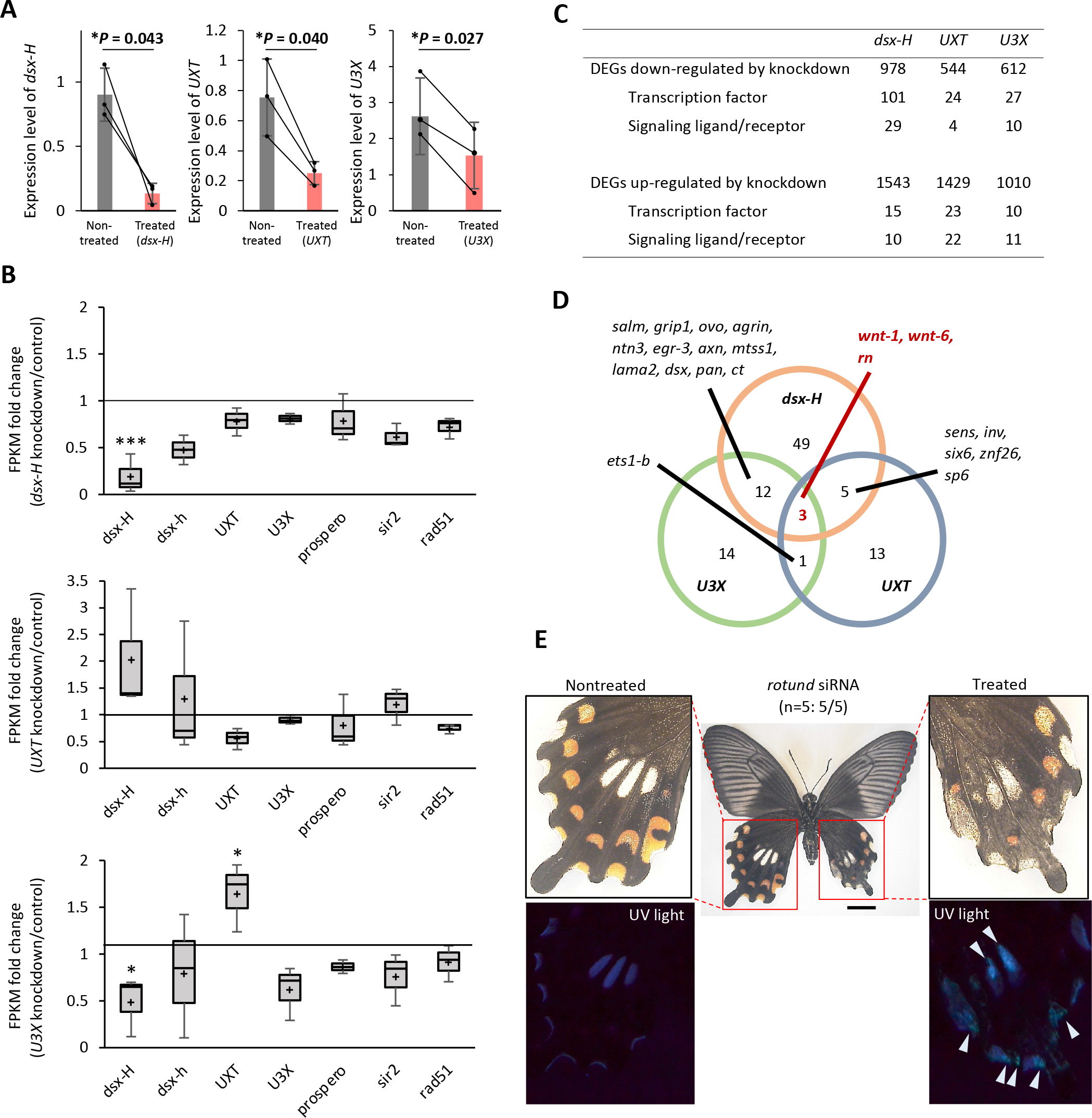
Comparison of gene expression levels between knockdown and control hindwings, and knockdown of *rotund* (*rn*) (**A**) Measurement of knockdown effect using RT-qPCR. We compared the expression levels of *dsx-H*, *UXT* and *U3X* between nontreated (grey bar) and treated hindwings (red bar) by RT-qPCR using *RpL3* as an internal control. Values and error bars denote the mean and standard deviation of three biological replicates. \**P* < 0.05 for paired t-test. (**B**) Fold Change of FPKM values (knockdown/control sides) of genes around *H* locus during knockdowns of *dsx-H*, *UXT* and *U3X*. FPKM fold changes of *dsx-H*, *dsx-h*, *UXT*, *U3X*, *prospero*, *sir2*, and *rad51* are shown. *rad51* is a gene adjacent to *sir2* (Fig. 1B), but its involvement in mimetic pattern formation has not been investigated to date. The value is 1 when the FPKM values of the siRNA treated and untreated sides are equal. Wald-test, paired; \*\*\**P*<0.001, \**P*<0.05. (**C**) The number of differentially expressed genes (DEGs) identified by *dsx-H*, *UXT* and *U3X* knockdowns. (**D**) Venn diagram depicting the abundance of DEGs (*P* < 0.05) for each comparison between three genes by untreated and siRNA-treated samples and shows only the number of transcription factors and signal factors. (**E**) Knockdown of *rn* in the hindwings of mimetic (*Hh*) females of *Papilio polytes*. siRNA was injected into the left pupal hindwing immediately after pupation and electroporated into the ventral side. Knockdown of *rn* changed the pale-yellow spots to produce UV fluorescence. UV fluorescence is not originally seen in mimetic females. White arrowheads represent the changed pale-yellow regions by knockdown. Scale bars, 1cm. Supplementary Fig. S12 show other replicates.

### The regulatory relationship and downstream genes of the three genes inside the HDR-*H*

Since all three genes in the HDR-*H*, *dsx-H*, *U3X*, and *UXT*, were found to be involved in the formation of mimetic patterns, we decided to examine their regulatory relationships and downstream genes. On the second day after injection (P2), siRNA un-injected hindwings (control) and injected hindwings (knockdown) were sampled for RNA extraction.

First, we confirmed siRNA injections of *dsx-H*, *UXT* and *U3X* reduced the expression levels of the target genes (Fig. 6A). We next examined the expression levels of genes inside and flanking the HDR (*dsx-H*, *dsx-h*, *UXT*, *U3X*, *prospero*, *sir2*, *rad51*) upon knockdown of *dsx-H*, *UXT*, and *U3X*. When *dsx-H* was knocked down, the expression of *dsx-H* was significantly decreased, but no significant expression changes were observed in other genes (Fig. 6B). Similarly, when *UXT* was knocked down, the expression of *UXT* tended to decrease (not statistically significant), but there was no significant effect on the expression of other genes (Fig. 6B). On the other hand, notably, when *U3X* was knocked down, in addition to the downward trend of *U3X* expression (not statistically significant), the expression of *dsx-H* was significantly decreased and the expression of *UXT* was significantly increased (Fig. 6B).

Comparing gene expressions in control and knockdown hindwings, about 500 to 1500 differentially expressed genes (DEGs) were extracted as genes whose expression was decreased or increased when each gene was knocked down (Fig. 6C).

We focused on the transcription factors and signaling factors whose expression is promoted by *dsx-H*, *UXT*, and *U3X* (Figs. S14 and S15), and found that *wnt1*, *wnt6*, and *rotund* (*rn*) were commonly down-regulated by knockdown of *dsx-H*, *UXT*, and *U3X* (Fig. 6D). *wnt1* and *wnt6* have been reported to be involved in the mimetic pattern formation (Iijima *et al*. 2019). When we knocked down *rn*, there was no characteristic change in the mimetic pattern, but there was an overall change in the color of the black, red, and pale-yellow regions, which seemed to become lighter (Figs. 6E and S16). When observed under UV irradiation in the *rn* knockdown wings, the UV fluorescence was observed in the pale-yellow spots (Figs. 6E and S16), where UV fluorescence is not observed usually in the mimetic form. From these observations, we consider that *rn* plays an important role in the pigment synthesis characteristic of the mimetic phenotype. Also, some of the transcription factors whose expression is reduced when *dsx-H* is knocked down include genes with color switching functions in other insects, such as *six6* (*optix*) and *bab2*. *optix* is involved in red spot formation in the wings of *Heliconius* butterflies and *bric a brac* (*bab*) is involved in abdominal pigmentation in *Drosophila* (Williams et al. 2008; Reed et al. 2011). It would be interesting to further examine these downstream genes in *P*. *polytes*.

Finally, comparing the expression of mimetic and non-mimetic females at P5 hindwings, 65 genes were extracted as upregulated in mimetic females and 47 genes as upregulated in non-mimetic females. Among the genes upregulated in mimetic females, 12 genes were found to be involved in some signaling pathways such as the *Wnt* signaling pathway (Table S5). On the other hand, only two of the upregulated genes in non-mimetic females were related to signaling pathways (Table S6).

## Discussion

In this paper, using the *in vivo* electroporation mediated RNAi method, we show that not only *dsx*, but also *UXT* and *U3X* in the inversion region for the *H*-allele, furthermore even outside flanking genes *prospero* and *sir2* are involved in the mimetic wing pattern formation in *P*. *polytes* (Figs. 4, 5 and 7A). The transcription factor *dsx* has been thought to function as a mimicry supergene as a single gene because it induces downstream genes to form the mimetic trait (Kunte et al. 2014; Baral et al. 2019). However, the present experiments indicate that multiple genes are involved in pattern formation. These genes are not downstream genes of *dsx-H* (Fig. S14), but are likely to function as members of the supergene. On the other hand, it is also important to note that we found that the expression of *U3X* may induce the expression of *dsx-H* (Fig. 6B). *U3X* is a long non-coding RNA not found in the *h*-allele and other genomic regions and is thought to have arisen specifically in HDR-*H* during evolution. *U3X* is located upstream of the transcription start site of *dsx-H*, and *U3X* may cis-regulates *dsx-H* expression. In *Daphnia magna*, long non-coding RNAs are also present upstream of *dsx* and regulate *dsx* function (Katoh et al. 2018), which indicates that further investigating more details of the regulatory mechanism of *U3X* expression are necessary. The knockdown of *U3X* did not necessarily cause a change from the mimetic to the non-mimetic patterns, but the RNA-seq results showed that *U3X* also repressed the expression of *UXT*, suggesting that the knockdown of *U3X* may have had the effect of increasing the gene expression of *UXT* (Figs. 4C and 6E). Knockdown of *UXT* switches the pattern to resemble the non-mimetic phenotype, including flattening of the upper part of the pale-yellow spots in the center of the hindwing (Fig. 4A). Importantly, the mosaic knockout of *UXT* in Crispr/Cas9 resulted in a similar phenotypic change (Fig. 4B), and the results were consistent between the two completely different experimental methods.

**Fig. 7.**
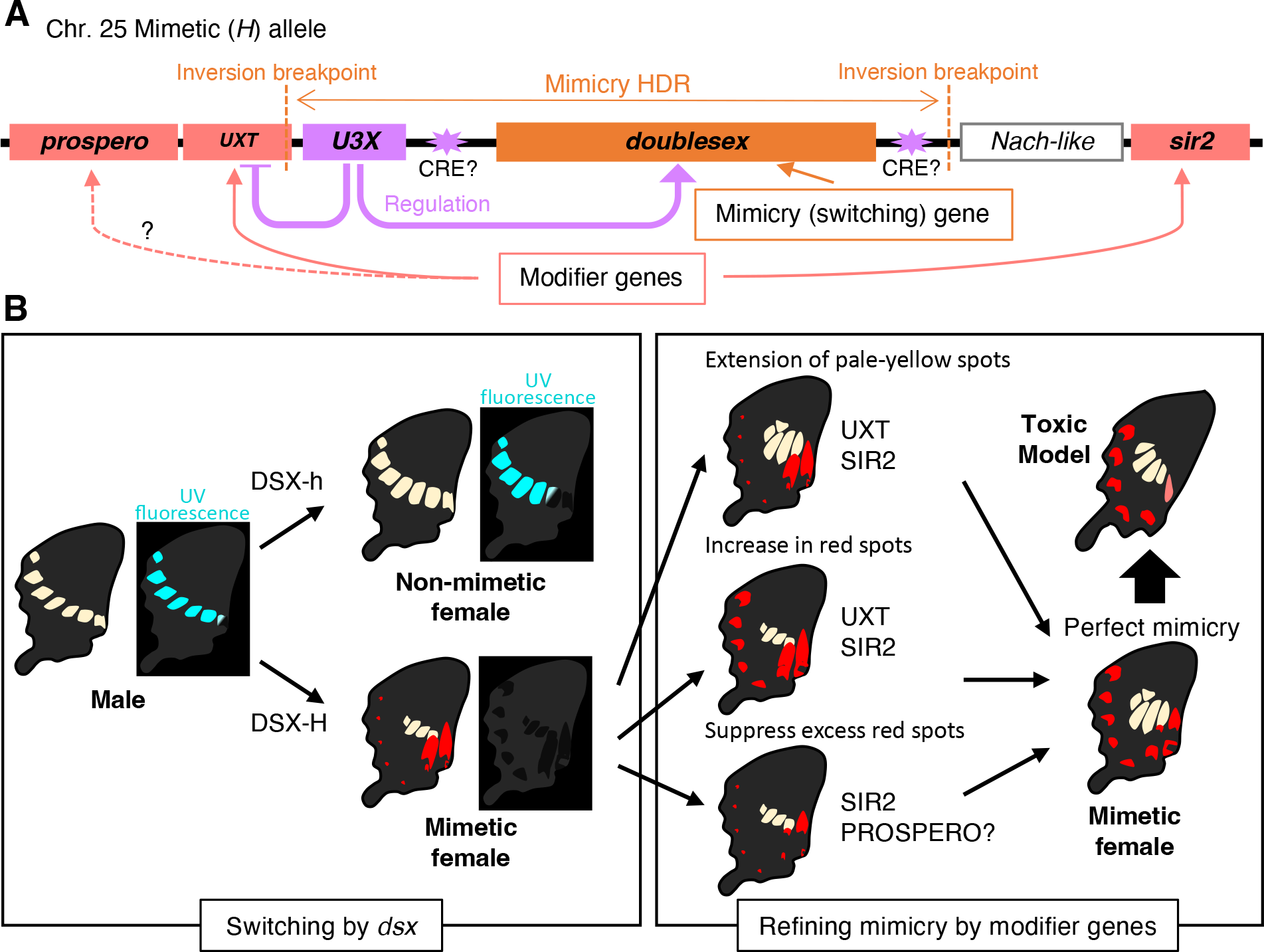
Model diagram of supergene structure and formation of mimetic color pattern by multiple genes in supergene of *Papilio polytes*. (**A**) The mimicry highly diversified region (HDR) on chromosome 25 contains three genes, *dsx*, *U3X*, and *UXT*, and neighboring genes such as *prospero*, *nach-like*, and *sir2*. It is hypothesized that *dsx* works as mimicry (switching) gene and that *UXT, sir2* and probably *prospero* functions as modifier genes. *U3X* upregulates the *dsx-H* expression and represses the *UXT* expression, and that there may be *cis*-regulatory element (CRE) which influences expressions of supergene members. (**B**) *dsx-h* switches from male to non-mimetic female and *dsx-H* switches from non-mimetic to mimetic female. The color pattern of the hindwings of mimetic females can be divided into three main parts. One is the pale-yellow spot, which is enlarged by the function of *UXT* and *sir2*. The second is the red spot on the outer edge of the hindwing, which is enlarged by *UXT* and *sir2*. The third is the red spot below the innermost pale-yellow spot, where the red spot is removed by the action of *sir2* and *prospero*. These modifier genes make the mimicry more like the model.

By functional analysis of *dsx-h*, which has been considered to have no specific function, it was shown that *dsx-h* induced non-mimetic patterns in females (Fig. 2B). This clearly indicates that the non-mimetic pattern in males is a default trait, and if *dsx-h* is expressed there, it becomes non-mimetic female, and if *dsx-H* is expressed there, it becomes mimetic female (Fig. 7B). This result suggests that *dsx-h* was originally involved in the regulation of sexual dimorphism in wing pattern, and the recombination was suppressed by an inversion, resulting in the differentiation of *dsx-H* and the evolution of female-limited Batesian mimicry. In butterflies, the evolution of female-limited polymorphism based on sexual dimorphism has been frequently hypothesized from evolutionary studies (Baral et al. 2019; Hopkins and Kopp 2021). Furthermore, functional analysis suggests that only isoform 3 of *dsx-H*, induces a mimetic pattern among the three female isoforms in this study (Fig. 3). The expression levels of each isoform were not significantly different (Fig. 1F), suggesting that the function of isoform 3 as a protein is important for the induction of mimetic traits, rather than the regulation of expression. Dsx is a transcription factor involved in sexual differentiation, and each isoform binds to a different response element, suggesting that the downstream gene network may change among three isoforms. Iijima et al. (2019) explored the downstream gene network of *dsx-H* for all isoforms, and it may be necessary to explore the downstream genes specific to isoform 3 of *dsx-H* for clarifying the mimicry mechanism in the future.

In recent years, many examples have been reported of supergenes in which complex adaptive phenotypes showing intraspecific polymorphism are regulated throughout a certain region of the chromosome (Gutiérrez-Valencia *et al*. 2021; Villoutreix *et al*. 2021), but this study is the first to investigate the functions of multiple genes in and flanking the HDR and to show that the gene cluster adjacent to *dsx* work as a supergene (Fig. 7A). *dsx-H* is thought to switch the phenotype from a non-mimetic to a mimic phenotype, and genes such as *UXT* and *sir2* are thought to make the mimetic phenotype more similar to the model (Fig. 7B). Because *dsx-H* changes not only the color pattern but also the pigment and scale microstructure specific to the mimetic form, genes such as *dsx-H* are called the mimicry gene, while those such as *UXT* and *sir2* are called modifier genes that are fine-tuned to improve mimicry (Charlesworth and Charlesworth 1975a; 1975b; Turner 1987; Charlesworth 2016). It is predicted that the mimicry gene evolved first, and modifier genes evolved later (Charlesworth and Charlesworth 1975a; 1975b; Turner 1987; Charlesworth 2016). We hypothesize about the evolution of the mimicry supergene in *P*. *polytes* as follows. First, inversion occurred around *dsx*, and then *dsx-H* and *U3X* originated, and mimicry females evolved, then *U3X* and *cis*-regulatory elements in the HDR may establish a regulatory mechanism for the expression of surrounding genes, and these genes may come to act as modifier genes (Fig. 7).

On the other hand, the results of expression analysis of each gene do not clearly indicate the regulatory relationships among genes in and flanking the mimicry HDR, and whether each gene is involved in the control of mimicry pattern formation (Fig. 1, C and D). In this study, all mRNA samples were prepared from the entire hindwing, and thus if a gene is expressed in a specific region (e.g., red spot region), it may not be possible to clearly judge the functional involvement of the gene in a mimetic pattern from the expression level. The only way to solve this problem is to compare the expression of each gene by *in situ* hybridization. In addition, we here compared gene expression levels at only three developmental timing: W (the first stage of the prepupa), P2, and P5. In order to obtain clear results, it is necessary to continuously compare gene expression levels at a wider range of time points. Furthermore, due to technical limitations, electroporation-mediated RNAi (siRNA injection) in the wing can only be performed immediately after pupation, which may not necessarily correspond to the time when each gene is functioning. In the case of *dsx-H* knockdown, it is noteworthy that the mimetic pattern is switched to the non-mimetic pattern even if RNAi is performed immediately after pupation (Nishikawa *et al*. 2015), suggesting that the fate of pattern formation is carried over at least to the early pupal stage. If RNAi can be applied to other stages of development, the functional role of each gene can be more clarified.

We would like to reconsider what is a supergene. Historically, it was assumed that multiple genes work together to produce more complex traits and to prevent recombination by placing genes adjacent to each other on the chromosome to avoid intermediate forms in the next generation, and such regions were defined as supergene (Fisher 1930; Darlington and Mather 1949; Ford 1965; Hamilton 1964; Dobzhansky 1970). In many of supergenes, chromosomal inversions are observed, and the structural diversity of multiple alleles is thought to be fixed by the inversions. And, Thompson and Jiggins (2014) defined supergene as ’a genetic architecture involving multiple linked functional genetic elements that allows switching between discrete, complex phenotypes maintained in a stable local polymorphism’. In the case of the female-limited polymorphic Batesian mimicry of *P*. *polytes* and its close relative, *P*. *memnon*, the whole genome sequence and GWAS showed that the causative region of the mimicry was a 150-kb region including *dsx* on chromosome 25 (Iijima *et al*. 2018). Both species have two types of low homology sequences (HDRs) corresponding to mimetic and non-mimetic alleles, but there is an inversion between the two alleles in *P*. *polytes*, but not in *P*. *memnon* (Iijima *et al*. 2018). It is not clear how sequence diversity arose and was maintained in *P*. *memnon*, but at least in *P*. *memnon*, the supergene cannot be defined in terms of the internal region of inversion. Then, it may be possible to define a supergene inside an HDR with low sequence homology, but is it possible to define a supergene including outer regions adjacent to an HDR with low sequence homology? This is an important question for understanding how we should think about the unit of the supergene and how the supergene has evolved.

Most supergene by an inversion contain more than a few dozen genes (some large supergenes contain more than 100 genes) (Gutiérrez-Valencia *et al*. 2021; Villoutreix *et al*. 2021). However, there has been no evidence that multiple genes belonging to the supergene are involved in complex adaptive traits. The fact that the mimicry supergene of *P*. *polytes* is only 130 kb in size and contains only three genes in the inversion region makes it more suitable than other supergenes for answering the above questions. In addition, it is a great advantage to be able to discuss it in comparison with the supergene of a related species, *P*. *memnon*, which does not have an inversion. Further investigation of gene function around the HDR, using multiple closely related species, will reveal more details about the function and evolution of the supergene.

Our present results suggest that the unit of the mimicry supergene can be defined to include at least the external neighboring genes. The results of comparative transcriptome analysis after knockdown of*dsx-H*, *U3X*, and *UXT* showed that the expressions of *sir2* and *prospero* were not the downstream genes of *dsx-H*, *U3X*, and *UXT*, suggesting that *sir2* and *prospero* expression is likely to be regulated by some *cis-*elements within HDR-*H*. The existence and location of *cis*-regulatory elements need to be investigated in the future, including possible epigenetic regulation of multiple genes in HDR-*H*. The significance of having related genes adjacent to each other on the chromosome should also be re-considered from the perspective of such expression regulation. For example, in the past, recombination sites may have been located further out and HDR dimorphism may have been more widespread, including adjacent *sir2* and *prospero*. In the process of evolution, *sir2* and *prospero* acquired functions involved in the pattern formation in addition to their original functions, and if these genes are able to work, they may not necessarily be in the recombination repression region. However, the regulation of their expressions may need to be affected by *cis*-regulatory elements inside the HDR-*H*.

## Acknowledgments

We thank Drs. T. Kojima and T. Iijima for helpful comments and experimental supports.

## Author contributions

HF conceived the study; SK, SY, YK, SS and KT conducted experiments; SK and HF wrote the paper. H F supervised this project. All authors reviewed the manuscript.

## Data and materials availability

The raw sequence data were deposited in DNA data bank of Japan (DDBJ). Accession information: transcriptome sequence accession ID, SAMD00000018646, SAMD00018647, SAMD00018649–SAMD00018657 (BioProject: PRJDB2955), SAMD00128718, SAMD00128715 (BioProject: PRJDB7141), SAMD00534475–SAMD00534484 and SAMD00535548–SAMD00535563 (BioProject: PRJDB14337).

