## Supplementary materials for "Functional unit and regulatory mechanisms of supergene in female-limited Batesian mimicry of *Papilio polytes*"

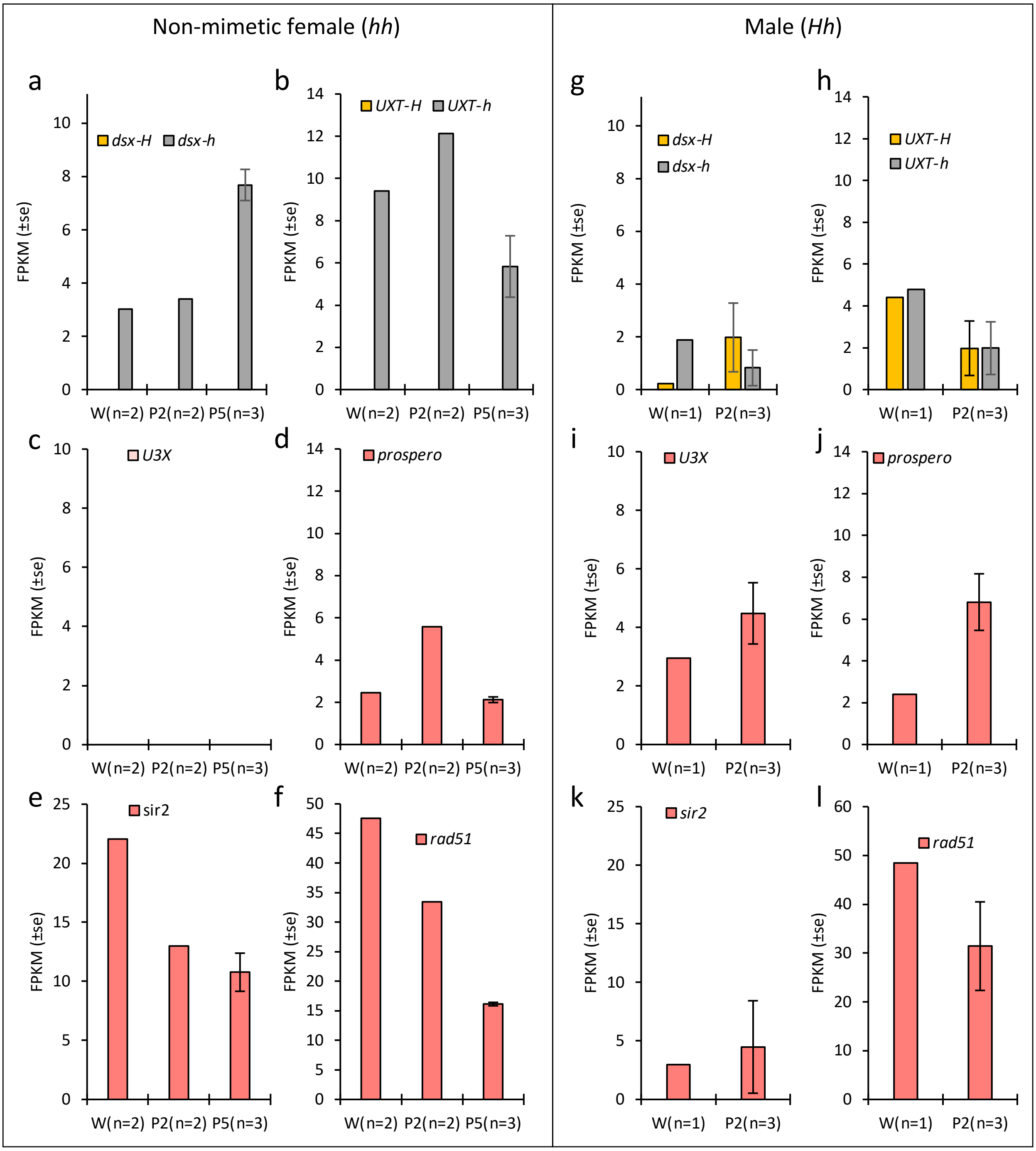


**Supplementary Figure S1.** Expression levels of genes within and flanking the HDR in hindwings of non-mimetic (*hh*) females and male (*Hh*) at the wandering stage (W) of the late last instar larvae, 2 day after pupation (P2) and 5 day after pupation (P5). The mean fragment per kilobase of transcript per million mapped reads (FPKM) values by RNA sequencing are shown with SE.

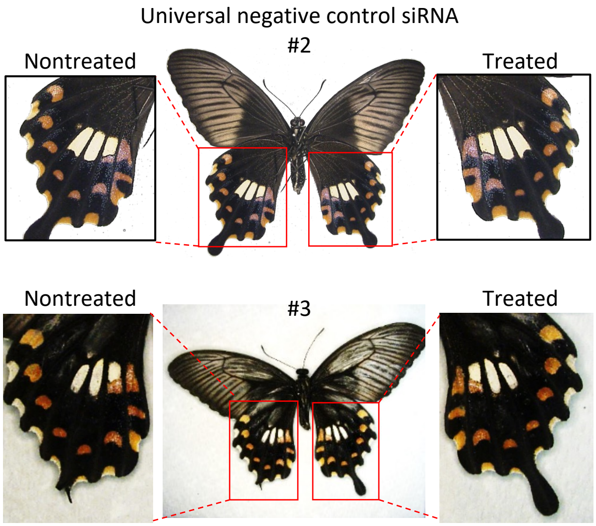


**Supplementary Figure S2.** A negative control using Universal Negative Control siRNA (Nippongene). No phenotypic effects were detected. Other replicates of Fig. 2A.

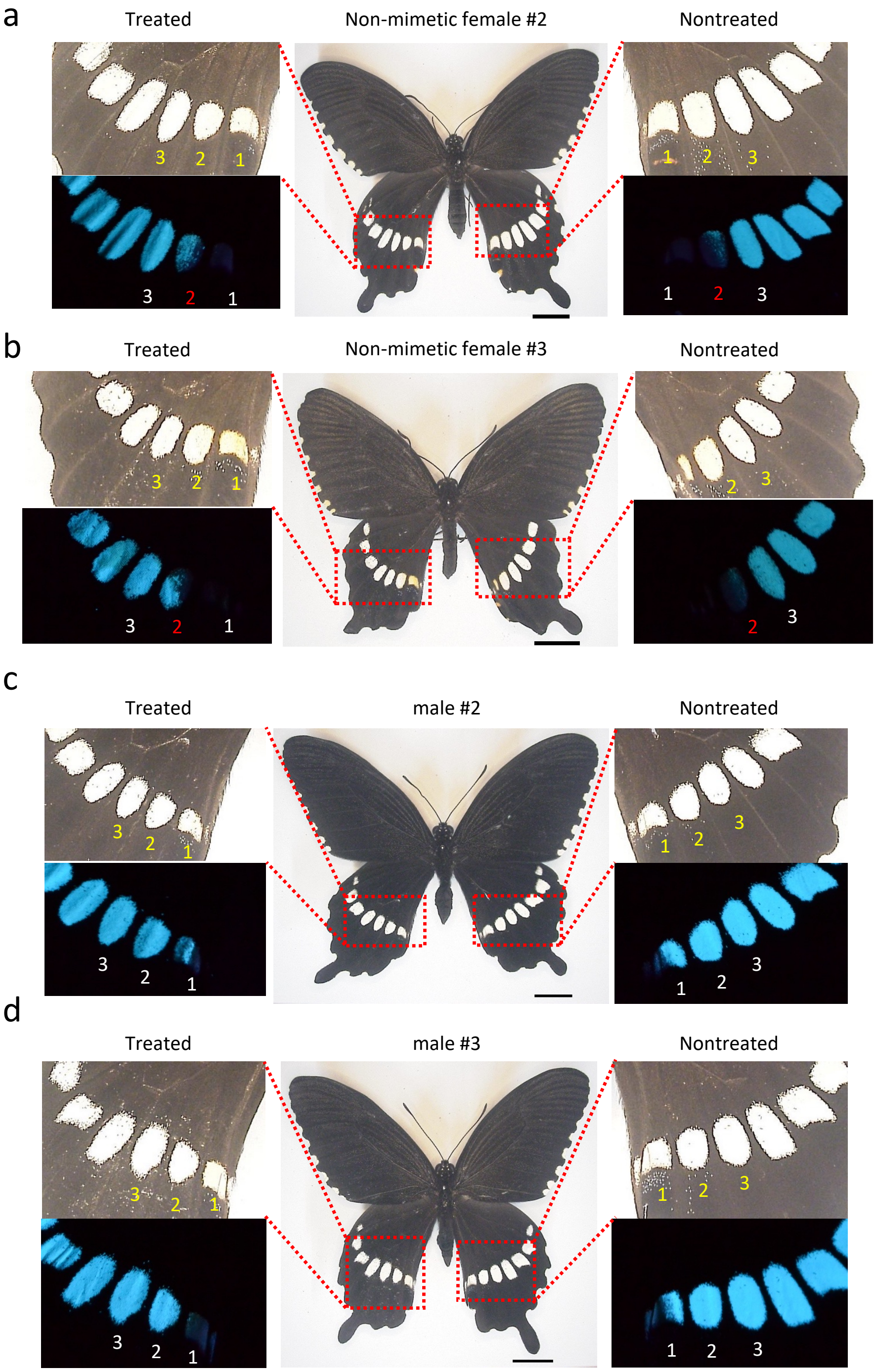


**Supplementary Figure S3.** Knockdown of *dsx* in the left hindwings of non-mimetic (*hh*) female (a,b) and male (c,d) of *Papilio polytes*. The siRNA targeting sequence common to all alleles and isoforms of *dsx* (*dsx-H*&*dsx-h* siRNA) was injected into the left pupal hindwing immediately after pupation and electroporated into the dorsal side. Other replicates of Fig. 2, B and C.

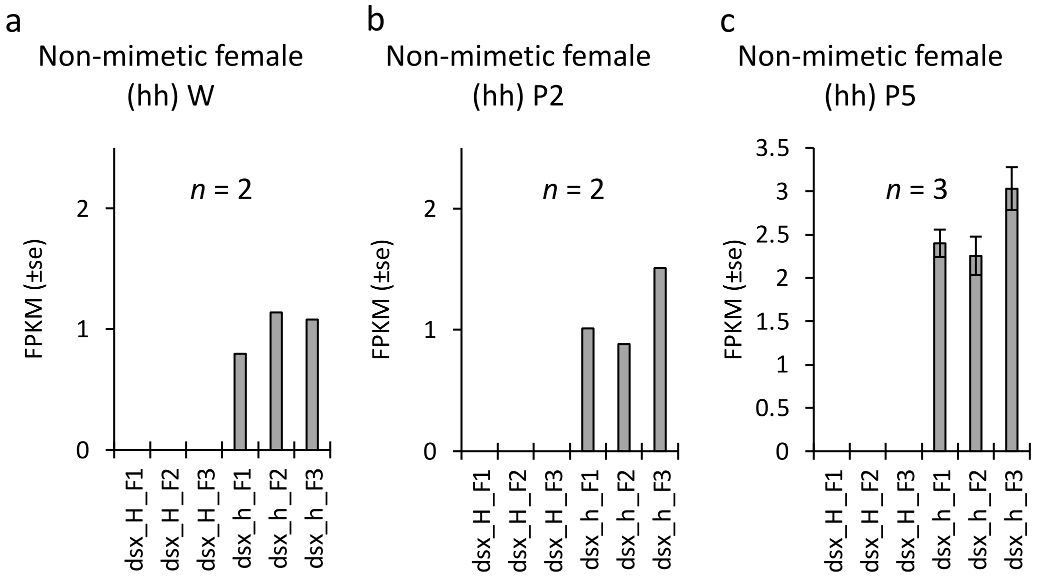


**Supplementary Figure S4.** Gene expression levels of each *dsx* isoforms in non-mimetic (*hh*) females in *Papilio polytes*. The mean fragment per kilobase of transcript per million mapped reads (FPKM) values by RNA sequencing at the wandering stage (W) of the late last instar larvae is (a), at 2 day after pupation (P2) is (b) and at 5 day after pupation (P5) is (c). Orange bars indicate the expression levels of dsx isoforms from mimetic (*H*) allele and gray bars indicate from non-mimetic (*h*) allele. F1, F2 and F3 means female isoform 1, 2 and 3, respectively. There was no statistically significant difference among isoforms.

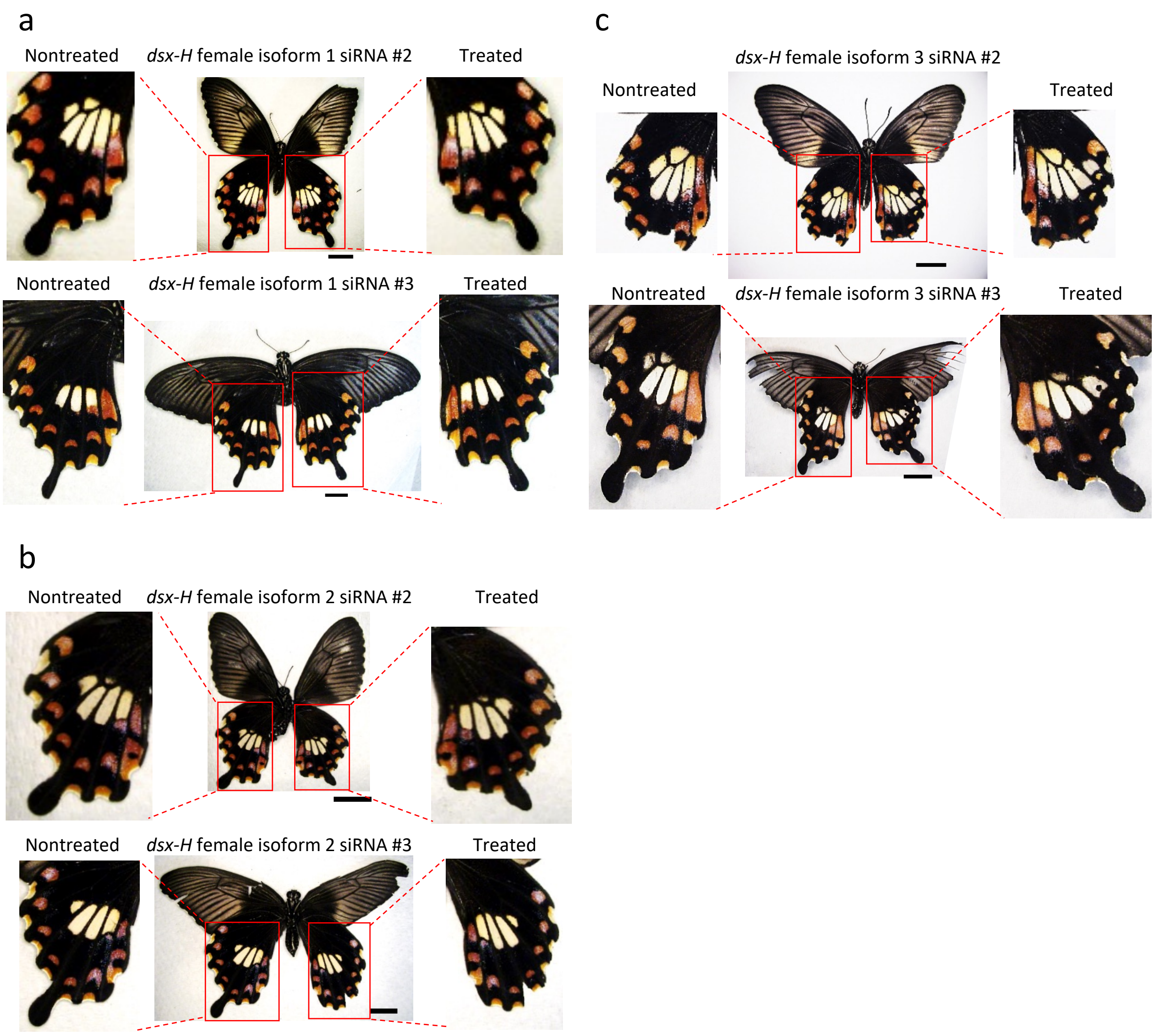


**Supplementary Figure S5.** Knockdown of dsx female isoform 1 (a), 2 (b) and 3 (c) in the hindwings of mimetic (*Hh*) females of *Papilio polytes*. Other replicates of Fig. 3.

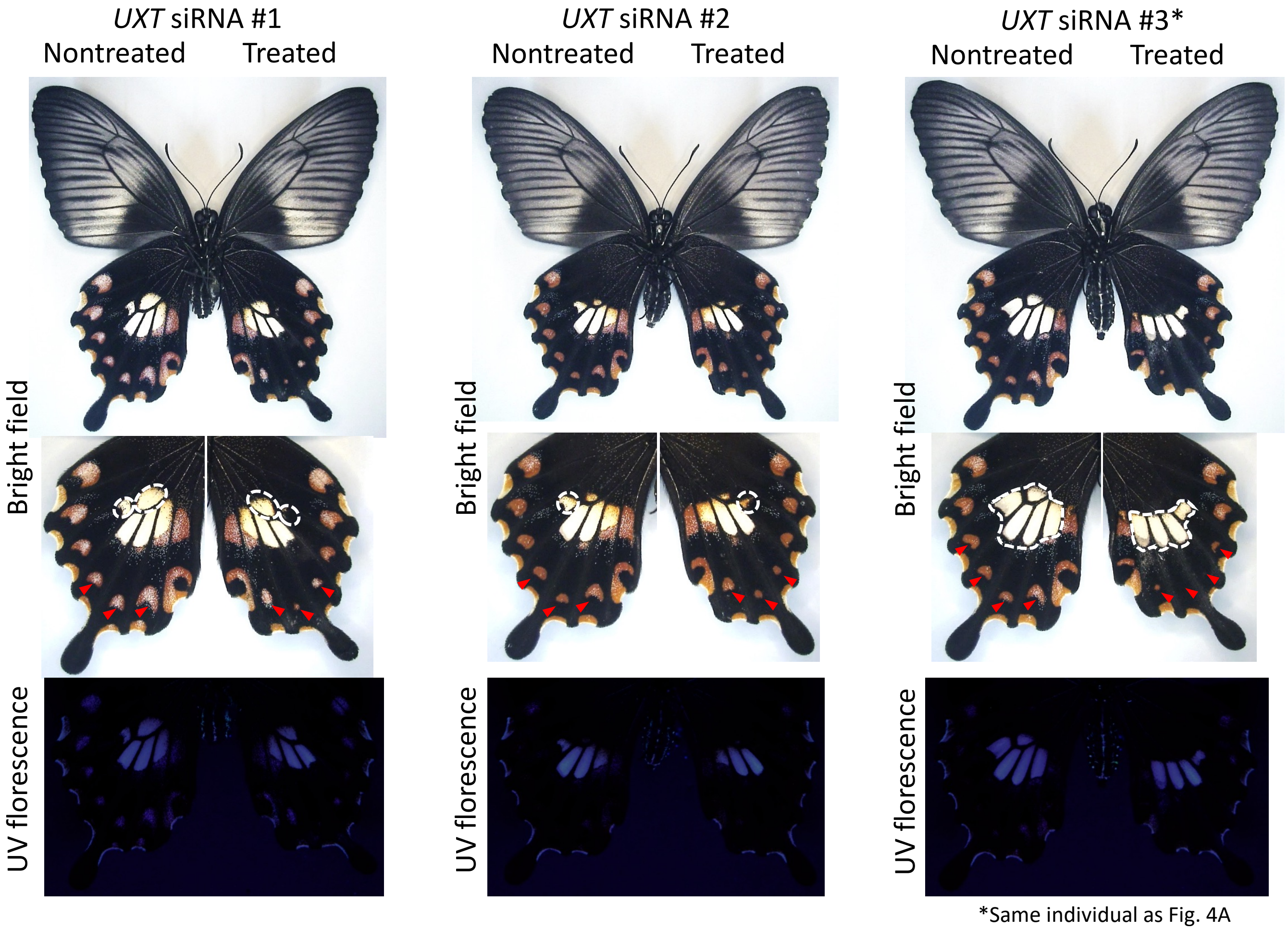


**Supplementary Figure S6.** Knockdown of *UXT* in the hindwings of mimetic (*Hh*) females. Other replicates of Fig. 4A (#3 is the same individual as Fig. 4A). Red arrowheads represent the changed red regions, and the white dotted line indicates the area where the pale yellow regions has changed.

a

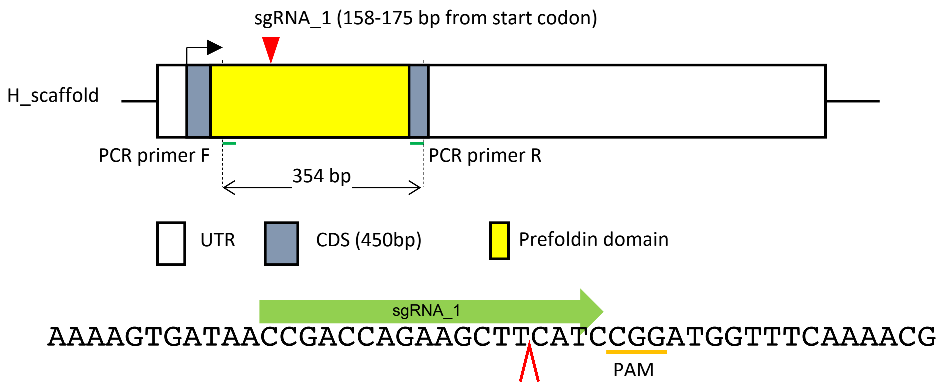


b

| | No. of injecting eggs | No. of hatching eggs | Hatching rates (%) | No. of 1-2 instars | No. of 3-5 instars | No. of pupa | No. of adult | Survival rates from larva to adult |
|---|---|---|---|---|---|---|---|---|
| plate1 | 54 | 22 | 40.74 | 14 | 8 | 7 | 5 | 22.72 |
| plate2 | 54 | 16 | 29.63 | 9 | 4 | 1 | 0 | 0.00 |
| plate3 | 54 | 20 | 37.04 | 16 | 7 | 6 | 4 | 20.00 |
| plate4 | 54 | 24 | 44.44 | 20 | 6 | 3 | 3 | 12.50 |
| plate5 | 54 | 16 | 29.63 | 16 | 8 | 5 | 4 | 25.00 |
| plate6 | 24 | 17 | 70.83 | 10 | 7 | 7 | 5 | 29.41 |
| total | 294 | 115 | 39.12 | 85 | 40 | 29 | 21* | 18.26 |

* Mimetic female = 8, nonmimetic female = 5, male = 8

**Supplementary Figure S7.** (a) Design of guide RNA in Crispr/Cas9 knockout experiment in *UXT*. PCR primers F and R show the primers for genotyping (Figure S8). (b) Summary of the number of eggs injected and the number of adults obtained in the Crispr/Cas9 experiment.

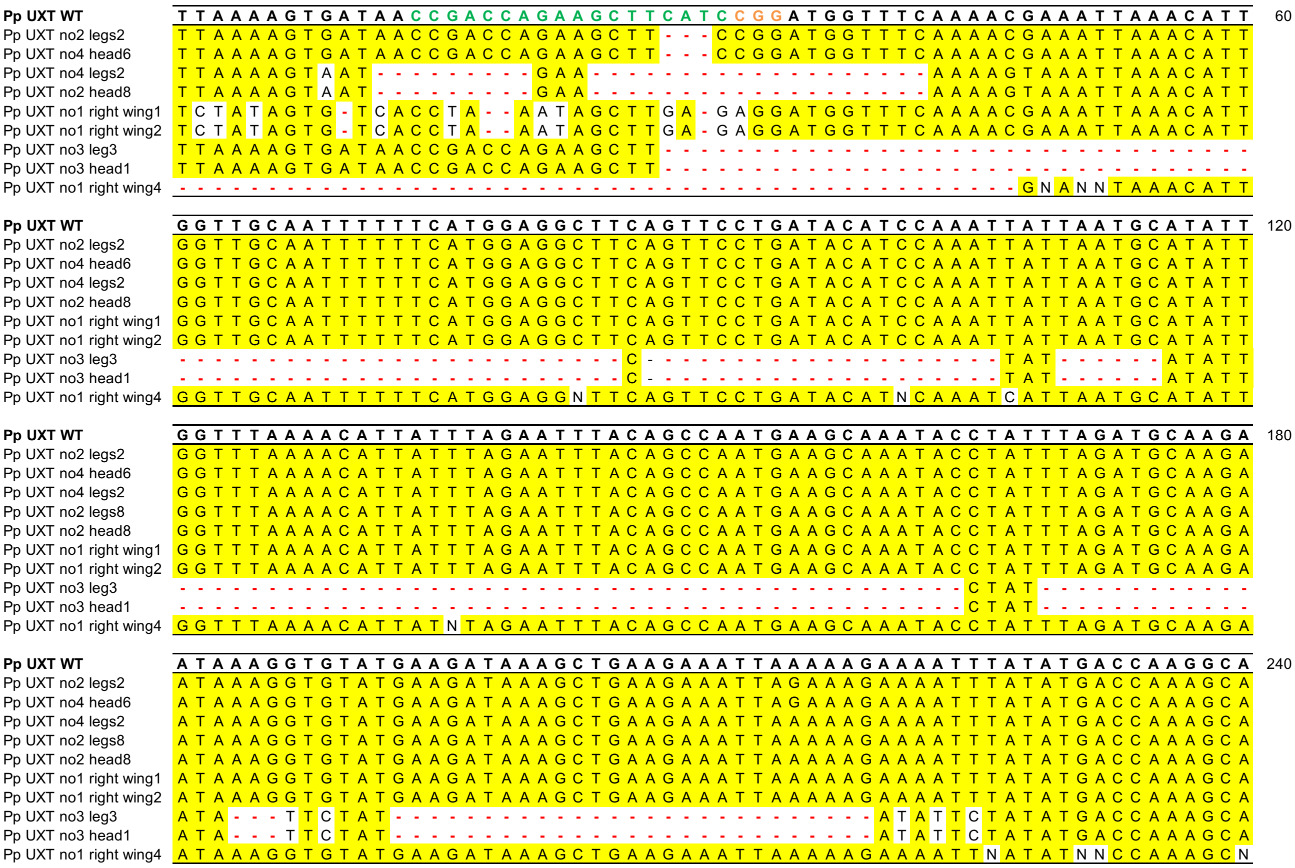


**Supplementary Figure S8.** CRISPR mutagenesis confirmed by Sanger sequencing. The top row shows the *UXT* sequence of *P. polytes*, the green sequence shows the target of sgRNA, and the orange shows the pam sequence.

(continue to the next page)

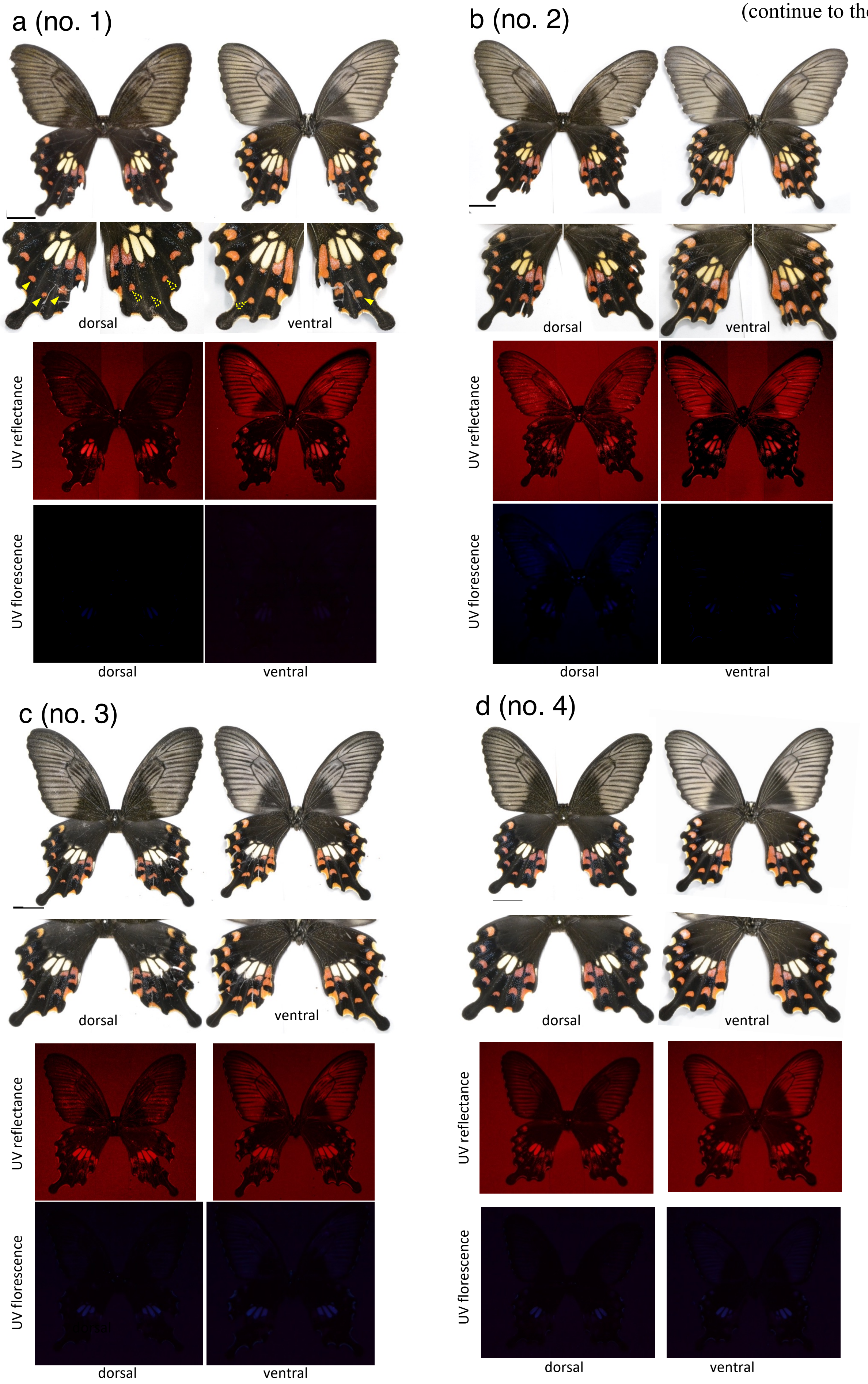


**Supplementary Figure S9.** Mosaic knockout of *UXT* by Crispr/Cas9. Dorsal and ventral views of eight individuals observed are shown. In individual number 1, 5, 6 and 8, arrowheads represent the changed red regions. The individual number 8 is the same individual shown in Fig. 4B. Scale bars, 1 cm. Other replicates of Fig. 4B.

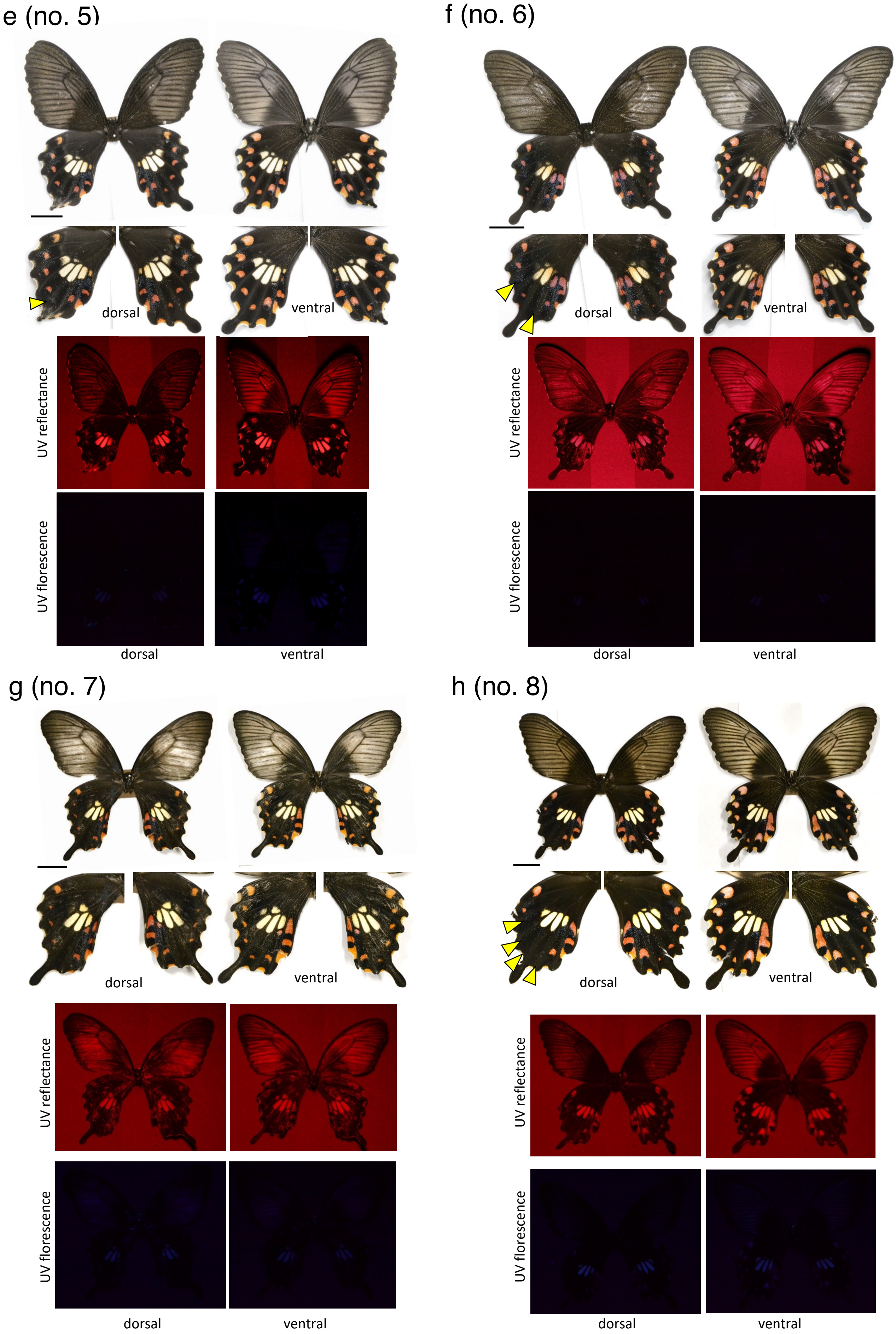

e (no. 5)
dorsal
ventral
UV reflectance
UV florescence
dorsal
ventral
f (no. 6)
dorsal
ventral
UV reflectance
UV florescence
dorsal
ventral
g (no. 7)
dorsal
ventral
UV reflectance
UV florescence
dorsal
ventral
h (no. 8)
dorsal
ventral
UV reflectance
UV florescence
dorsal
ventral

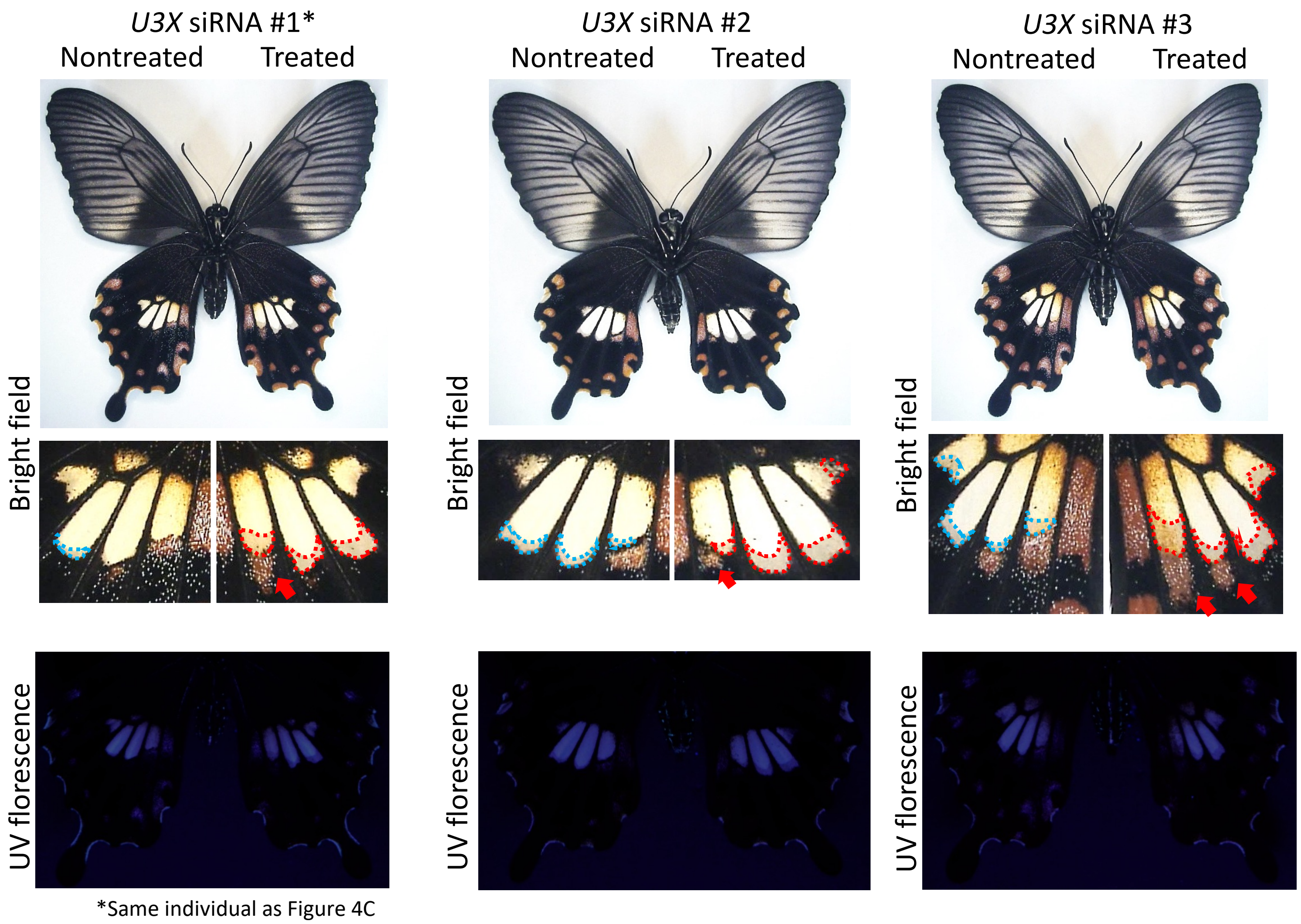


**Supplementary Figure S10.** Knockdown of *U3X* in the hindwings of mimetic (*Hh*) females. In magnified views of the pale yellow regions of *U3X* knockdown , the red arrow indicates the area where the red spot has expanded, and the red dotted line indicates the area where the pale yellow spot has extended. Other replicates of Fig. 4C (#1 is the same individual as Fig. 4C.).

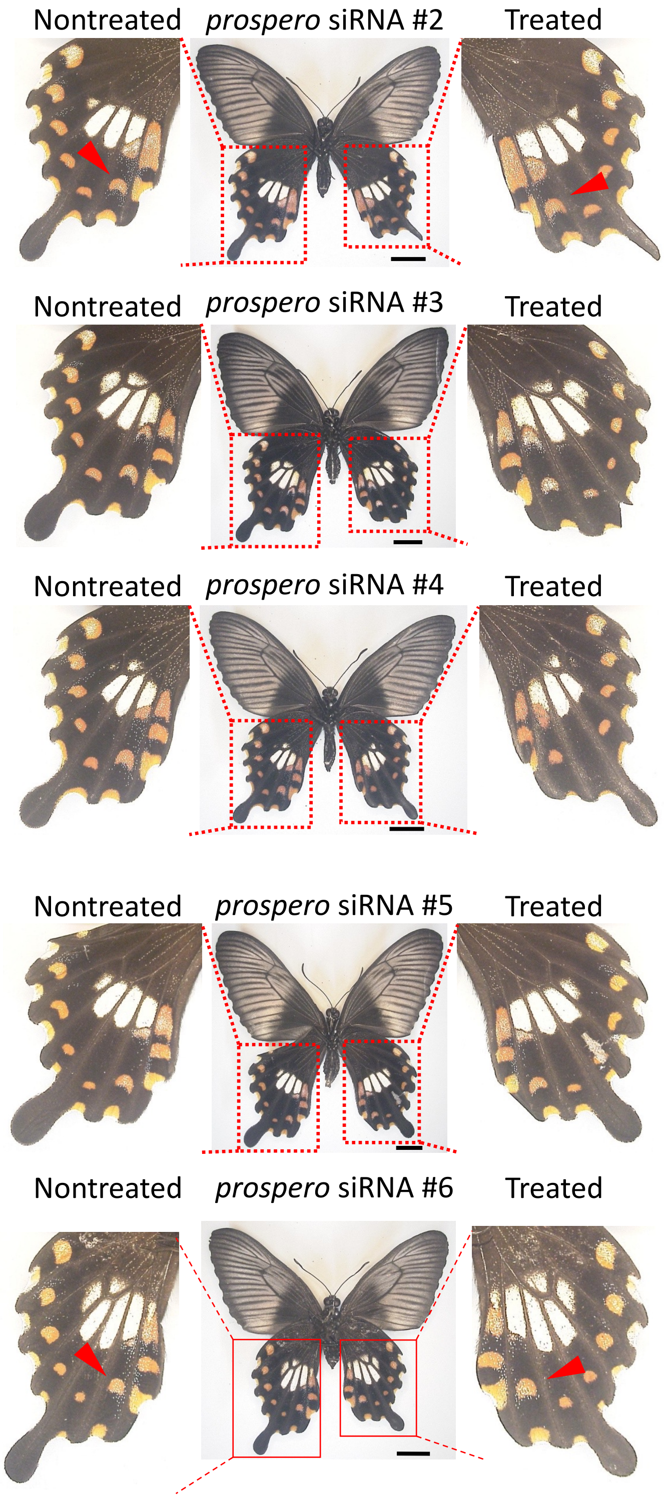


**Supplementary Figure S11.** Knockdown of *prospero* in the hindwings of mimetic (*Hh*) females. As in Fig. 5A, in #2 and #6, the red spot indicated by the red arrowhead is faintly enlarged on the treated hindwing. Other replicates of Fig. 5A.

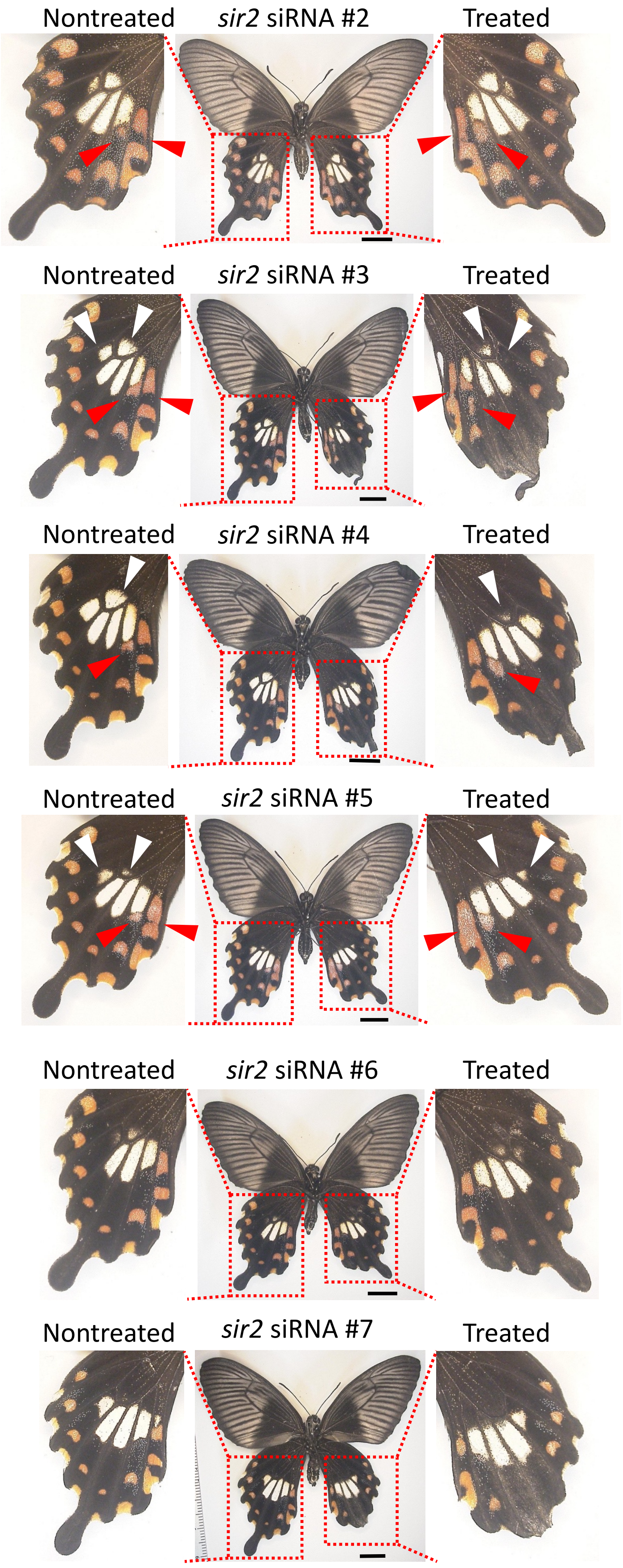


**Supplementary Figure S12.** Knockdown of *sir2* in the hindwings of mimetic (*Hh*) females. Other replicates of Fig. 5B. Red and white arrowheads represent the changed red and pale-yellow regions, respectively.

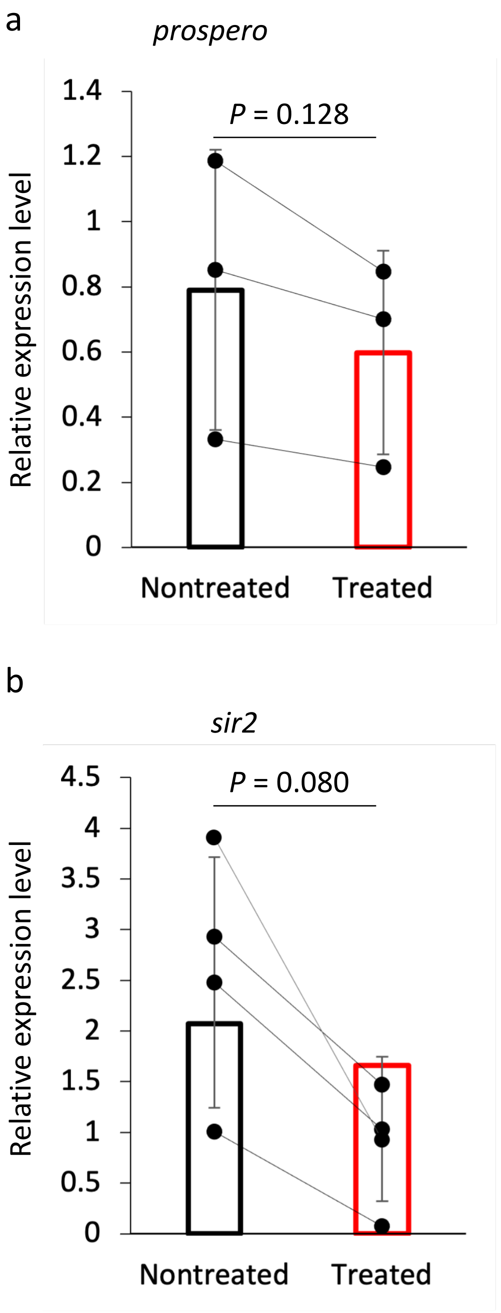


**Supplementary Figure S13.** Measurement of knockdown effect using RT-qPCR. We compared the expression levels of *prospero* (a) and *sir2* (b) between untreated (white bar) and treated hindwings (red bar) by RT-qPCR using *RpL3* as an internal control. Values and error bars denote the mean and standard deviation of three biological replicates. *P* values were obtained by one-tailed paired t-test.

a

| Name | Official Gene Symbol | Uniprot ID | $Log_2FC$ | *P*-value |
|---|---|---|---|---|
| **DSX** | LOC106110500 | P23023 | 4.45675 | 0.0108380750 |
| HMOC | LOC106100350 | P22810 | 4.27549 | 0.0000463000 |
| MECOM | LOC106110943 | Q03112 | 3.61564 | 0.0258608690 |
| SOX21 | LOC106101338 | Q811W0 | 3.48508 | 0.0000529000 |
| DICH | LOC106101336 | Q24533 | 3.27739 | 0.0035838870 |
| SIX6 | LOC106103966 | O95475 | 2.83884 | 0.0080575900 |
| LBX2 | LOC106102952 | Q804R0 | 2.15327 | 0.0000170000 |
| GFI1 | LOC106099801 | Q9N658 | 1.9022 | 0.004301761 |
| AWH | LOC106102621 | Q8IRC7 | 1.80563 | 0.00062436 |
| SP9 | LOC106109488 | Q0VA40 | 1.75262 | 0.002509899 |
| TBX20 | LOC106099923 | Q3SA46 | 1.64003 | 0.0000027 |
| ODD | LOC106105176 | P23803 | 1.61629 | 0.008206448 |
| FD4 | LOC106099254 | P32028 | 1.51677 | 0.000509026 |
| BAB2 | LOC106105236 | Q9W0K4 | 1.47143 | 0.000458556 |
| EMS | LOC106105889 | P18488 | 1.44825 | 0.003622841 |
| FSA6 | LOC106103511 | A0A0E4AZF8 | 1.42681 | 0.0262287 |
| AHR | LOC106110299 | P41738 | 1.41849 | 0.017579811 |
| SP3 | LOC106109077 | Q90WR8 | 1.37218 | 0.009419911 |
| **RN** | LOC106111359 | Q9VI93 | 1.28911 | 0.01272464 |
| SALM | LOC106099579 | P39770 | 1.28646 | 0.003775342 |
| SLP2 | LOC106111539 | P32031 | 1.27369 | 0.018884271 |
| SOB | LOC106105175 | Q9VQS7 | 1.26472 | 0.000569916 |
| AP2E | LOC106108726 | Q2T9K2 | 1.24444 | 0.02308945 |
| LHX5 | LOC106107010 | Q9H2C1 | 1.19358 | 0.003148396 |
| CI | LOC106111406 | P19538 | 1.18878 | 0.000957467 |
| ZN182 | LOC106110817 | P17025 | 1.13764 | 0.003696544 |
| ZN497 | LOC106101137 | Q6ZNH5 | 1.07692 | 0.034082211 |
| UNC4 | LOC106106876 | O77215 | 1.04558 | 0.004324119 |
| TRH | LOC106104251 | Q24119 | 1.04547 | 0.026487951 |
| HMIN | LOC106101873 | P27610 | 1.04394 | 0.018330703 |
| ZNF57 | LOC106102639 | Q68EA5 | 1.00517 | 0.008836051 |
| CRBL2 | LOC106105347 | Q642H2 | 0.99423 | 0.003428941 |
| CUT | LOC106111596 | P10180 | 0.98809 | 0.001874532 |
| HTH | LOC106101088 | O46339 | 0.98422 | 0.006361462 |
| BOWEL | LOC106105174 | Q9VQU9 | 0.92879 | 0.023964653 |
| PURB | LOC106111297 | Q6PHK6 | 0.90919 | 0.025784516 |
| ABRU | LOC106101269 | Q24174 | 0.87095 | 0.003625336 |
| OVO | LOC106100378 | P51521 | 0.85323 | 0.036968437 |
| P53 | LOC106109054 | P79734 | 0.84172 | 0.006516223 |
| SCAL | LOC106104996 | P30052 | 0.81112 | 0.022769686 |
| BC11A | LOC106101611 | Q9H165 | 0.79043 | 0.03416502 |
| HAIR | LOC106107661 | P14003 | 0.76552 | 0.004745104 |
| EGR3 | LOC106103066 | P43300 | 0.75955 | 0.040394836 |
| ZNF26 | LOC106105951 | P17031 | 0.75487 | 0.027343338 |
| AL | LOC106099537 | Q06453 | 0.73129 | 0.022670832 |
| ZN648 | LOC106101930 | Q5T619 | 0.72847 | 0.012496932 |
| MEF2 | LOC106111879 | P40791 | 0.7228 | 0.00998445 |
| PANG1 | LOC106111393 | P91943 | 0.70176 | 0.033990271 |
| TRX | LOC106104170 | Q24742 | 0.69468 | 0.018742801 |
| ZN792 | LOC106107921 | Q3KQV3 | 0.6448 | 0.019024689 |
| MYB | LOC106110522 | P01103 | 0.62841 | 0.023424575 |
| ZN423 | LOC106111049 | A1L1R6 | 0.59202 | 0.030765891 |
| DA | LOC106101365 | P11420 | 0.5257 | 0.04353915 |

b

| Name | Official Gene Symbol | Uniprot ID | $Log_2FC$ | *P*-value |
|---|---|---|---|---|
| LAMA2 | LOC106109953 | Q60675 | 1.90801 | 0.0000711 |
| AGRIN | LOC106100673 | P31696 | 1.38281 | 0.007098057 |
| PTK7 | LOC106108697 | B3MH43 | 1.25805 | 0.001511162 |
| GRIP1 | LOC106100211 | P97879 | 1.25193 | 0.019375575 |
| **WNT6** | LOC106099738 | P22727 | 1.24225 | 0.0000666 |
| MTSS1 | LOC106109943 | O43312 | 1.21592 | 0.000375392 |
| SYNE1 | LOC106111486 | Q6ZWR6 | 1.15591 | 0.007647198 |
| NET3 | LOC106101675 | Q90923 | 1.15064 | 0.000866058 |
| TGFR1 | LOC106099197 | Q64729 | 1.05795 | 0.00057184 |
| AXN | LOC106107355 | Q9V407 | 1.03366 | 0.00388509 |
| **WNT1** | LOC106099686 | P49340 | 1.03263 | 0.000360695 |
| PRTG | LOC106102724 | Q2EY15 | 1.02864 | 0.009759818 |
| NMDE2 | LOC106109676 | Q5R1P3 | 0.82011 | 0.040585245 |
| ATRN | LOC106106136 | O75882 | 0.8034 | 0.015550742 |
| NLGN3 | LOC106099319 | Q8BYM5 | 0.66227 | 0.02892004 |
| ERBIN | LOC106108590 | Q96RT1 | 0.66215 | 0.034623186 |

$-1 < Log_2FC \leq 0$
$-2 < Log_2FC \leq -1$
$-3 < Log_2FC \leq -2$
$-4 < Log_2FC \leq -3$
$Log_2FC \leq -4$

$0.01 \leq p < 0.05$
$0.001 \leq p < 0.01$
$p < 0.001$

**Supplementary Figure S14.** DEGs of transcription factors (a) and signaling factors (b) down-regulated by *dsx-H* knockdown. Sequences of DEGs obtained by RNA-seq were blasted with Uniprot, and transcription factors and signal factors were extracted using GO terms of the top hit Uniprot ID. In case multiple isoforms of the same gene were included, only the isoform with the lowest $Log_2FC$ (Log2-fold change value = Log2 siRNA-treated FPKM–Log2 untreated FPKM) is shown, and Name is the protein name of the top hit Uniprot ID.

a

| Name | Official Gene Symbol | Uniprot ID | Log$_2$FC | *P*-value |
|---|---|---|---|---|
| BARH1 | LOC106107800 | Q24255 | -2.679 | 0.04043 |
| SIX6 | LOC106103966 | O95475 | -1.6967 | 0.03596 |
| SP9 | LOC106109488 | Q0VA40 | -1.6001 | 0.008 |
| GFI1 | LOC106099801 | Q9N658 | -1.4969 | 0.00973 |
| GLIS3 | LOC106111309 | Q6XP49 | -1.4714 | 0.01423 |
| ZN679 | LOC106111510 | Q8IYX0 | -1.3806 | 0.01517 |
| **RN** | LOC106111359 | Q9VI93 | -1.1748 | 0.04265 |
| HMIN | LOC106101873 | P27610 | -1.0504 | 0.01992 |
| ZNF45 | LOC106101236 | Q02386 | -0.8835 | 0.02833 |
| ZF316 | LOC106101613 | Q6PGE4 | -0.8284 | 0.00482 |
| ETS1B | LOC106101086 | P18756 | -0.7943 | 0.0098 |
| ZNF26 | LOC106105951 | P17031 | -0.7464 | 0.01526 |
| ZN431 | LOC106100240 | E9QAG8 | -0.671 | 0.0375 |
| ZN676 | LOC106105927 | Q8N7Q3 | -0.6375 | 0.03193 |
| AL | LOC106099903 | Q06453 | -0.624 | 0.03937 |
| E2F4 | LOC106109225 | Q16254 | -0.5866 | 0.04339 |
| ZN454 | LOC106108144 | Q8N9F8 | -0.584 | 0.02974 |
| LOLA1 | LOC106106644 | P42283 | -0.4937 | 0.04617 |

b

| Name | Official Gene Symbol | Uniprot ID | Log$_2$FC | *P*-value |
|---|---|---|---|---|
| SPON2 | LOC106105716 | Q9WV75 | -2.0535 | 0.01542 |
| **WNT1** | LOC106099686 | P49340 | -1.38 | 0.02905 |
| **WNT6** | LOC106099738 | P22727 | -1.2292 | 0.00919 |
| PICK1 | LOC106110046 | Q9EP80 | -1.0243 | 0.04986 |

$-1<Log_2FC\leq 0$
$-2<Log_2FC\leq -1$
$-3<Log_2FC\leq -2$
$-4<Log_2FC\leq -3$
$Log_2FC\leq -4$

$0.01 \leq p<0.05$
$0.001\leq p<0.01$
$p<0.001$

c

| Name | Official Gene Symbol | Uniprot ID | Log$_2$FC | *P*-value |
|---|---|---|---|---|
| **DSX** | LOC106110500 | P23023 | 1.46124 | 0.0473 |
| GLIS2 | LOC106107952 | Q7K0S9 | 1.06861 | 0.0277 |
| PAX2 | LOC106100458 | P32114 | 0.96373 | 0.0160 |
| CUT | LOC106111596 | P10180 | 0.78737 | 0.0026 |
| PANG1 | LOC106111393 | P91943 | 0.76656 | 0.0119 |
| ESMB | LOC106110754 | Q01069 | 0.75679 | 0.0118 |
| FRU | LOC106104066 | Q8IN81 | 0.66947 | 0.0216 |
| EGL13 | LOC106105518 | Q23045 | 0.63449 | 0.0337 |
| EGR3 | LOC106103066 | P43300 | 0.59276 | 0.0425 |
| ETS1B | LOC106101086 | P18756 | 0.58116 | 0.0178 |
| DLL | LOC106101899 | P20009 | 0.52863 | 0.0312 |
| **RN** | LOC106111359 | Q9VI93 | 0.52546 | 0.0121 |
| TGIF2 | LOC106102596 | Q8C0Y1 | 0.52458 | 0.0300 |
| SALL1 | LOC106099579 | Q9NSC2 | 0.48446 | 0.0284 |
| OVO | LOC106100378 | P51521 | 0.43677 | 0.0168 |
| SP3 | LOC106101378 | Q90WR8 | 0.43336 | 0.0443 |
| ELF1 | LOC106100333 | P13002 | 0.37986 | 0.0180 |
| ZNF84 | LOC106107173 | P51523 | 0.35197 | 0.0401 |
| NFXL1 | LOC106108291 | A0JMY5 | 0.33536 | 0.0441 |
| TCRG1 | LOC106101251 | Q8CGF7 | 0.30729 | 0.0332 |

d

| Name | Official Gene Symbol | Uniprot ID | Log$_2$FC | *P*-value |
|---|---|---|---|---|
| GRIP1 | LOC106100211 | P97879 | 0.97899 | 0.0455 |
| NET3 | LOC106101675 | Q90923 | 0.7392 | 0.0097 |
| MTSS1 | LOC106109943 | O43312 | 0.67824 | 0.0128 |
| AGRIN | LOC106100673 | P31696 | 0.65683 | 0.0432 |
| **WNT6** | LOC106099738 | P22727 | 0.61202 | 0.0022 |
| LAMA2 | LOC106109953 | Q60675 | 0.55741 | 0.0082 |
| **WNT1** | LOC106099686 | P49340 | 0.55419 | 0.0253 |
| AXN | LOC106107355 | Q9V407 | 0.49305 | 0.0249 |
| SRC42 | LOC106102457 | Q9V9J3 | 0.48769 | 0.0490 |
| ADA10 | LOC106103498 | O14672 | 0.35938 | 0.0210 |

**Supplementary Figure S15.** DEGs of transcription factors and signaling factors down-regulated by *UXT* (a,b) and *U3X* (c,d) knockdown. Among the DEGs obtained by knockdown of *UXT*, transcription factors are shown in (a) and signal factors are shown in (b). Similarly, the transcription factor and signal factor of *U3X* are shown in (c) and (d), respectively.

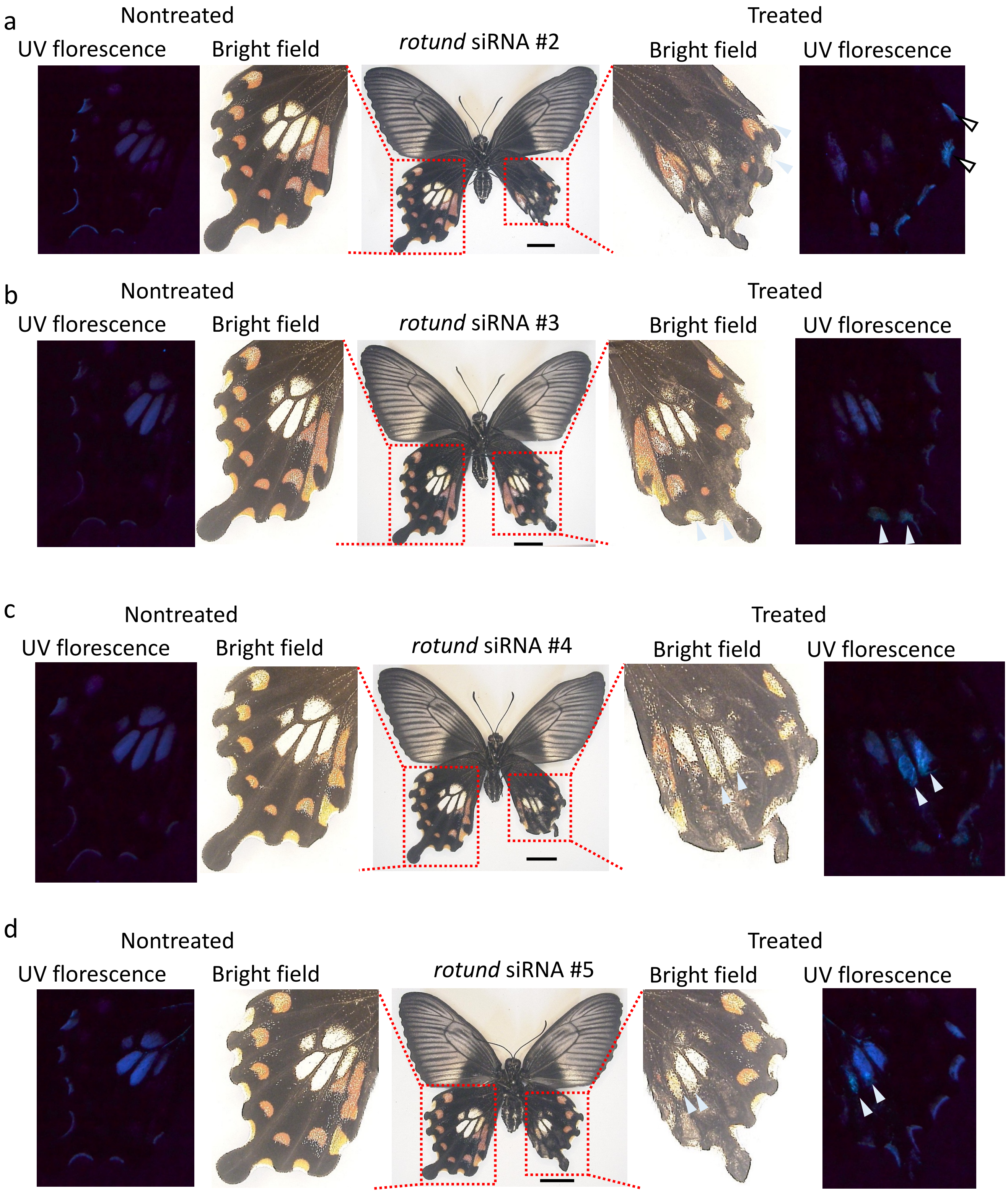


**Supplementary Figure S16.** Knockdown of *rotund* (*rn*) in the hindwings of mimetic (*Hh*) females. Other replicates of Fig. 6E 11. In UV fluorescence, white arrowheads represent the changed pale yellow regions by knockdown. Knockdown of *rn* changed the pale yellow spots to produce UV fluorescence like males. Scale bars, 1cm.

**Supplementary Table S1. Summary of RNA-seq data for analyzing expression levels in hindwing.**

| sex | genotype | stage* | Total reads | Accession number | Reference |
|---|---|---|---|---|---|
| female | Hh (mimetic) | W | 32662442 | SAMD00018653 | Nishikawa et al. 2015 |
| female | Hh (mimetic) | W | 43756504 | SAMD00018654 | Nishikawa et al. 2015 |
| female | Hh (mimetic) | W | 32073800 | SAMD00018655 | Nishikawa et al. 2015 |
| female | Hh (mimetic) | P2 | 44802896 | SAMD00018646 | Nishikawa et al. 2015 |
| female | Hh (mimetic) | P2 | 48917672 | SAMD00018647 | Nishikawa et al. 2015 |
| female | Hh (mimetic) | P2 | 34191856 | SAMD00018649 | Nishikawa et al. 2015 |
| female | Hh (mimetic) | P2 | 37108690 | SAMD00018650 | Nishikawa et al. 2015 |
| female | Hh (mimetic) | P5 | 57845316 | SAMD00534475 | this study |
| female | Hh (mimetic) | P5 | 42352556 | SAMD00534476 | this study |
| female | Hh (mimetic) | P5 | 52407784 | SAMD00534477 | this study |
| female | hh (non-mimetic) | W | 44210098 | SAMD00018651 | Nishikawa et al. 2015 |
| female | hh (non-mimetic) | W | 49523076 | SAMD00018652 | Nishikawa et al. 2015 |
| female | hh (non-mimetic) | P2 | 47905432 | SAMD00534478 | this study |
| female | hh (non-mimetic) | P2 | 46441150 | SAMD00534479 | this study |
| female | hh (non-mimetic) | P5 | 51941714 | SAMD00534480 | this study |
| female | hh (non-mimetic) | P5 | 52537276 | SAMD00534481 | this study |
| female | hh (non-mimetic) | P5 | 48761016 | SAMD00534482 | this study |
| male | Hh | W | 32114136 | SAMD00018657 | Nishikawa et al. 2015 |
| male | Hh | P2 | 54632630 | SAMD00534483 | this study |
| male | Hh | P2 | 47742400 | SAMD00534484 | this study |
| male | Hh | P2 | 52810550 | SAMD00018656 | Nishikawa et al. 2015 |

*W: 5th instar larvae (wandering), P2: 2 days after pupation, P5: 5 days after pupation

**Supplementary Table S2. Summary of RNA-seq data for DEGs between control and RNAi hind wings.**

| Sample type | sex | genotype | stage* | Total reads | Accession number |
|---|---|---|---|---|---|
| dsx-H_control1 | female | *HH* | P2 | 53406060 | SAMD00535549 |
| dsx-H_knockdown1 | female | *HH* | P2 | 67393442 | SAMD00535548 |
| dsx-H_control2 | female | *Hh* | P2 | 71828378 | SAMD00128715 |
| dsx-H_knockdown2 | female | *Hh* | P2 | 77618118 | SAMD00128718 |
| dsx-H_control3 | female | *Hh* | P2 | 47225584 | SAMD00535551 |
| dsx-H_knockdown3 | female | *Hh* | P2 | 42890920 | SAMD00535550 |
| UXT_control1 | female | *Hh* | P2 | 53548310 | SAMD00535553 |
| UXT_knockdown1 | female | *Hh* | P2 | 56542592 | SAMD00535552 |
| UXT_control2 | female | *Hh* | P2 | 47523810 | SAMD00535555 |
| UXT_knockdown2 | female | *Hh* | P2 | 56463070 | SAMD00535554 |
| UXT_control3 | female | *Hh* | P2 | 59193152 | SAMD00535557 |
| UXT_knockdown3 | female | *Hh* | P2 | 63099692 | SAMD00535556 |
| U3X_control1 | female | *Hh* | P2 | 46621574 | SAMD00535559 |
| U3X_knockdown1 | female | *Hh* | P2 | 64156422 | SAMD00535558 |
| U3X_control2 | female | *Hh* | P2 | 57740102 | SAMD00535561 |
| U3X_knockdown2 | female | *Hh* | P2 | 52851868 | SAMD00535560 |
| U3X_control3 | female | *Hh* | P2 | 52290574 | SAMD00535563 |
| U3X_knockdown3 | female | *Hh* | P2 | 59202374 | SAMD00535562 |

### Supplementary Table S3. List of qPCR primer

| Target gene | qPCR primer name | Formard (5'–3') | Reverse (5'–3') |
|---|---|---|---|
| *dsx* female isoform1 | dsx_F1 | GATAGATGAAGGAAAGCTCATCGTG | CCACCTTCGTCCGGTCATG |
| *dsx* female isoform2 | dsx_F2 | GGGAACTCGACACGCCAGTAT | TCATTCGCAAAACAACTTCA |
| *dsx* female isoform3 | dsx_F3 | GATAGATGAAGGAAAGCTCATCGTG | ATTTCGCTCAGCATTTTCTGGCG |
| *U3X* | Pp_U3X_qPCR | CCTGACATCTACTAAATGCTTCATGG | CAGCACCAGCTCGTGTTGCTTG |
| *UXT-H* | qPpUXT_mimetic_spec_F | CATTTTCGGAATGATGGATGCGATA | TAGCGGTATCAACATTTGGTAGAGA |
| *UXT-h* | qPpUXT_nonmimetic_spec_F | GACGCGATGTTTTTAGCTTTTTACA | TAGCGGTATCAACATTTGGTAGAGA |
| *dsx-H* | Pp_dsx_H_spec_qPCR | gctgcaactcaccacgcagcgtcaca | ccgcgctcggagtcgacggaggt |
| *dsx-h* | Pp_dsx_h_spec_qPCR | gctgcaacttaccacgcggcgcaact | ccgagctcgaagtcgacgggggc |
| *sir2* | Pp_Sir2_qPCR_1 | TTGTAGGAAACAATACTCTTTGGAA | GGCTTGATAATGCCTGGACA |
| *prospero* | Pp_prospero_qPCR_1 | GAGGTGCCACCCAACTTCAG | CGCGGAAGAATTCCCGTAA |
| *RpL3** | Pp_rpl3_qPCR-F2 | CACAAAGGGCAAGGGATAC | ACAAGCTACTTTACGCAGAC |

*used as an internal control

### Supplementary Table S4. Lists of siRNA

| siRNA target gene | siRNA name | siRNA Target sequence | Sense (5'–3') | Antisense (5'–3') |
|---|---|---|---|---|
| *dsx-H* female isoform 1 | Pp_dsxH_F1_A | AAGGTGGAGAAATTCGAAAAATA | GGUGGAGAAAUUCGAAAAAUA | UUUUUCGAAUUUCUCCACCUU |
| *dsx-H* female isoform 2 | Pp_dsxH_F2_A | TCGACACGCCAGTATGGACTTTA | GACACGCCAGUAUGGACUUUA | AAGUCCAUACUGGCGUGUCGA |
| *dsx-H* female isoform 3 | Pp_dsxH_F3_A | GTGTAGTATCGTCTTCTATGAAG | GUAGUAUCGUCUUCUAUGAAG | UCAUAGAAGACGAUACUACAC |
| *dsx-H* | Pp_dsx_mimetic | TTGTCGCAACCACCGGTTGAAGG | GUCGCAACCACCGGUUGAAGG | UUCAACCGGUGGUUGCGACAA |
| *dsx-H&dsx-h* | Pp_dsx_common | TTGGTGGAGAACTGTCACAGACT | GGUGGAGAACUGUCACAGACU | UCUGUGACAGUUCUCCACCAA |
| *UXT* | Pp_UXT_A | AAGGTGTATGAAGATAAAGCTGA | GGUGUAUGAAGAUAAAGCUGA | AGCUUUAUCUUCAUACACCUU |
| *U3X* | Pp_U3X_A | AAGAAACAACAAAATTACCATAT | GAAACAACAAAAUUACCAUAU | AUGGUAAUUUUGUUGUUUCUU |
| *sir2* | Pp_Sir2_C | AACCAACAATTTCACATTATTTC | CCAACAAUUUCACAUUAUUUC | AAUAAUGUGAAAUUGUUGGUU |
| *sir2* | Pp_Pmem_Sir2_D | TCCGTCATTACACACAGAATATT | CGUCAUUACACACAGAAUAUU | UAUUCUGUGUGUAAUGACGGA |
| *prospero* | Pp_prospero_A | AAGAACAGTTAGCTGAAATGAAA | GAACAGUUAGCUGAAAUGAAA | UCAUUUCAGCUAACUGUUCUU |
| *prospero* | Pp_prospero_C | AACAACAACGAGCCTAAATTAAA | CAACAACGAGCCUAAAUUAAA | UAAUUUAGGCUCGUUGUUGUU |
| *rotund* | Pp_ZNF_rotund_A | GAGCACATTCCTAAACACAAAGA | GCACAUUCCUAAACACAAAGA | UUUGUGUUUAGGAAUGUGCUC |

**Supplementary Table S5. DEGs upregulated in the hindwings of mimetic females (*Hh*) at 5 day after pupation (P5) compared to non-mimetic females (*hh*).** Sequences of DEGs obtained by RNA-seq were blasted with Uniprot, and transcription factors and signal factors were searched using GO terms ("DNA-binding Transcription factor activity" [GO:0003700], "DNA-binding transcription factor activity, RNA polymerase II-specific" [GO:0000981], "signaling receptor binding" [GO:0005102], "signaling receptor activity" [GO:0038023]) of the top hit Uniprot ID. In case multiple isoforms of the same gene were included, only the isoform with the lowest $Log_2FC$ (Log2-fold change value = Log2 siRNA-treated FPKM–Log2 untreated FPKM) is shown, and Name is the protein name of the top hit Uniprot ID. Since only two transcription factors and one signaling factor were found (highlighted in bold), we also searched for GO terms containing "signaling pathways" and listed their representative functions in the GO column.

| *Papilio polytes* Gene ID | Uniprot ID | Log2FC | P-value | Name | GO |
|---|---|---|---|---|---|
| XM_013292375.1 | P23023 | -8.028901 | 1.99E-06 | DSX_DROME | **Transcription factor** |
| XM_013278880.1 | A0JNC4 | -6.283797 | 0.0159084 | ELOV7_BOVIN | |
| XM_013289657.1 | Q9Y115 | -5.617943 | 0.0414717 | UN93L_DROME | |
| XM_013293168.1 | Q5RHR6 | -4.960954 | 0.0004459 | BROMI_DANRE | |
| XM_013289324.1 | Q10126 | -4.194233 | 1.84E-21 | YSM6_CAEEL | |
| XM_013293435.1 | Q62770 | -3.277414 | 0.0208379 | UN13C_RAT | |
| XM_013283047.1 | P15122 | -3.220498 | 0.0363557 | ALDR_RABIT | |
| XR_001225879.1 | Q9HH35 | -3.186314 | 0.03473 | FLPA_METWO | |
| XM_013283036.1 | Q9VER6 | -3.028281 | 0.0280421 | MODSP_DROME | Toll signaling pathway |
| XM_013278742.1 | Q9NQX1 | -2.956486 | 0.0133877 | PRDM5_HUMAN | **Transcription factor** |
| XM_013288907.1 | P55112 | -2.840364 | 0.0001654 | NAS4_CAEEL | |
| XM_013292531.1 | Q93126 | -2.813686 | 0.0035115 | GPR9_AMPAM | |
| XM_013281013.1 | Q9NL89 | -2.475945 | 4.22E-06 | BGBP_BOMMO | cell surface pattern recognition receptor signaling pathway |
| XM_013282109.1 | Q9ILI6 | -2.331282 | 0.0397911 | NS1A_TASV2 | |
| XM_013281764.1 | P38621 | -2.237384 | 0.0280421 | ZN12_MICSA | |
| XM_013293332.1 | P82147 | -2.186865 | 0.0024826 | L2EFL_DROME | |
| XM_013284803.1 | Q7ZV90 | -2.174838 | 0.0003456 | PIF1_DANRE | |
| XM_013282531.1 | A8I9E8 | -2.146214 | 0.0059134 | CFA45_CHLRE | |
| XM_013292524.1 | Q1EHB4 | -2.109849 | 0.0422167 | SC5AC_HUMAN | |
| XM_013278023.1 | Q9BTX7 | -1.963251 | 0.0163264 | TTPAL_HUMAN | |
| XM_013292079.1 | E2AX35 | -1.903661 | 0.0001654 | PROH4_CAMFO | neuropeptide signaling pathway |
| XM_013284791.1 | Q86MW9 | -1.885181 | 0.0233423 | SINA_SCHMA | |
| XM_013290665.1 | Q63ZT8 | -1.766709 | 0.0253732 | AL1L1_XENTR | |
| XM_013283469.1 | Q5NVG8 | -1.61198 | 0.0206612 | WASF1_PONAB | |
| XM_013292792.1 | Q9UHJ6 | -1.477074 | 0.03473 | SHPK_HUMAN | |
| XM_013281497.1 | P21674 | -1.344644 | 0.0084348 | FST_RAT | BMP signaling pathway |
| XM_013291468.1 | Q9QY36 | -1.315578 | 5.23E-05 | NAA10_MOUSE | |
| XM_013293424.1 | Q25490 | -1.287657 | 0.0066073 | APLP_MANSE | Wnt signaling pathway |
| XM_013287565.1 | Q32NZ6 | -1.286312 | 7.40E-07 | TMC5_MOUSE | |
| XM_013293116.1 | B4F6U4 | -1.277509 | 0.0084654 | PRD10_XENTR | |
| XM_013285557.1 | Q6GMR7 | -1.273569 | 0.000499 | FAAH2_HUMAN | |

| | | | | | |
|---|---|---|---|---|---|
| XR_001225479.1 | Q5D189 | -1.16555 | 0.0206996 | TYSY_BACNA | |
| XM_013282178.1 | P70478 | -1.130181 | 0.0036855 | APC_RAT | Wnt signaling pathway |
| XM_013294132.1 | Q9QUR6 | -1.114853 | 9.21E-06 | PPCE_MOUSE | |
| XM_013285687.1 | Q9W440 | -1.110978 | 0.048895 | THEM6_DROME | |
| XM_013281861.1 | P16568 | -1.10538 | 0.0468203 | BICD_DROME | |
| XM_013291878.1 | P56188 | -1.100689 | 0.0014197 | RPE_HELPY | |
| XM_013289041.1 | O45599 | -1.051108 | 0.0073054 | CBD1_CAEEL | |
| XM_013280429.1 | Q9V5N8 | -1.028348 | 1.69E-05 | STAN_DROME | cell surface receptor signaling pathway |
| XM_013285297.1 | Q7PHR1 | -0.948362 | 0.0280421 | KIF1A_ANOGA | |
| XM_013281987.1 | Q9W1A4 | -0.935817 | 0.0014197 | TAMO_DROME | |
| XM_013290545.1 | Q9ULD9 | -0.899649 | 0.0363557 | ZN608_HUMAN | |
| XM_013289108.1 | Q9I8X3 | -0.76056 | 0.0011665 | FGFR3_DANRE | transmembrane receptor protein tyrosine kinase signaling pathway |
| XM_013280973.1 | O88407 | -0.750667 | 0.0299537 | LFG2_RAT | negative regulation of apoptotic signaling pathway |
| XM_013283234.1 | Q5H8C4 | -0.747697 | 0.0160064 | VP13A_MOUSE | |
| XM_013280435.1 | A0A0R4IES7 | -0.714689 | 0.049824 | K1109_DANRE | |
| XM_013288394.1 | Q664K8 | -0.714598 | 0.0085942 | HSLO_YERPS | |
| XM_013280607.1 | P43304 | -0.708153 | 0.0035115 | GPDM_HUMAN | |
| XM_013277766.1 | P20241 | -0.691435 | 0.0280421 | NRG_DROME | |
| XM_013292770.1 | O15118 | -0.683172 | 0.0237362 | NPC1_HUMAN | **Signaling factor** |
| XM_013288992.1 | F1M3J4 | -0.682091 | 0.0006897 | MRP4_RAT | bile acid signaling pathway |
| XM_013293712.1 | P48809 | -0.67119 | 0.0301453 | RB27C_DROME | |
| XM_013288785.1 | Q9VLT5 | -0.666925 | 0.0248156 | POE_DROME | |
| XM_013293162.1 | Q14624 | -0.65939 | 0.0106412 | ITIH4_HUMAN | |
| XM_013290786.1 | Q95Q62 | -0.65428 | 0.0127745 | IP3KH_CAEEL | |
| XM_013277707.1 | Q03161 | -0.651129 | 0.0499764 | YMY9_YEAST | |
| XM_013283867.1 | Q8IWK6 | -0.635408 | 0.0463328 | AGRA3_HUMAN | cell surface receptor signaling pathway |
| XM_013293461.1 | Q811L6 | -0.626496 | 0.0346593 | MAST4_MOUSE | |
| XM_013293375.1 | Q9BLC5 | -0.613787 | 0.0460297 | HSP83_BOMMO | |
| XM_013281601.1 | P54356 | -0.602187 | 0.03473 | TSG_DROME | BMP signaling pathway |
| XM_013280987.1 | Q8BKX6 | -0.569993 | 0.0299537 | SMG1_MOUSE | |
| XM_013291108.1 | Q80TR8 | -0.568632 | 0.0328461 | DCAF1_MOUSE | |
| XM_013289542.1 | Q9QUR8 | -0.54338 | 0.0490491 | SEM7A_MOUSE | |
| XM_013289978.1 | Q00963 | -0.542764 | 0.0441267 | SPTCB_DROME | |
| XM_013278069.1 | Q66I79 | -0.518673 | 0.0454497 | SELK_DANRE | |

**Supplementary Table S6. DEGs upregulated in the hindwings of non-mimetic females (*hh*) at 5 day after pupation (P5) compared to mimetic females (*Hh*).** Sequences of DEGs obtained by RNA-seq were blasted with Uniprot, and transcription factors and signal factors were searched using GO terms ("DNA-binding Transcription factor activity" [GO:0003700], "DNA-binding transcription factor activity, RNA polymerase II-specific" [GO:0000981], "signaling receptor binding" [GO:0005102], "signaling receptor activity" [GO:0038023]) of the top hit Uniprot ID. In case multiple isoforms of the same gene were included, only the isoform with the highest $Log_2FC$ (Log2-fold change value = Log2 siRNA-treated FPKM–Log2 untreated FPKM) is shown, and Name is the protein name of the top hit Uniprot ID. Since no transcription factors and signaling factor was found, we also searched for GO terms containing "signaling pathways" and listed their representative functions in the GO column.

| *Papilio polytes* Gene ID | Uniprot ID | Log2FC | P-value | Name | GO |
|---|---|---|---|---|---|
| XM_013291377.1 | Q964T2 | 7.1006701 | 0.0185687 | CP9E2_BLAGE | |
| XM_013287918.1 | P26305 | 6.5611432 | 0.0035115 | LPSBP_PERAM | |
| XM_013293804.1 | P30432 | 5.8118372 | 0.0007167 | FUR2_DROME | |
| XM_013288243.1 | P08793 | 5.5220865 | 0.0005628 | XDH_CALVI | |
| XM_013279897.1 | A6H639 | 4.7158468 | 0.0059134 | DRC5_MOUSE | |
| XM_013278088.1 | P49098 | 4.3054509 | 0.0059443 | CYB5_TOBAC | |
| XM_013283708.1 | Q2NAK3 | 3.7767182 | 0.0103113 | RSMI_ERYLH | |
| XM_013285508.1 | Q06343 | 3.1956578 | 0.0002783 | BJSB2_TRINI | |
| XM_013293416.1 | B0WC46 | 2.9702478 | 0.0175462 | TRET1_CULQU | |
| XM_013292332.1 | A3KN33 | 2.9400823 | 0.0217315 | EGFLA_BOVIN | |
| XM_013281630.1 | Q3UTY6 | 2.8036817 | 0.0371378 | THSD4_MOUSE | |
| XM_013279400.1 | Q5R4U0 | 2.7971337 | 0.0195759 | CAH10_PONAB | |
| XM_013291941.1 | O01761 | 2.7908971 | 6.16E-07 | UNC89_CAEEL | |
| XM_013289082.1 | Q99943 | 2.7236901 | 0.0422167 | PLCA_HUMAN | cytokine-mediated signaling pathway |
| XM_013284531.1 | Q8R4G8 | 2.6366831 | 0.039754 | KCTD1_RAT | |
| XM_013286195.1 | Q95031 | 2.3973961 | 0.0271391 | CP6B6_HELAM | |
| XM_013281804.1 | Q9D110 | 2.3095503 | 0.0059134 | MTHFS_MOUSE | |
| XM_013293532.1 | A0A2R8QCI3 | 2.2331306 | 0.0257821 | DAPLE_DANRE | Wnt signaling pathway |
| XM_013286322.1 | P15145 | 1.9778471 | 0.0379987 | AMPN_PIG | |
| XM_013281867.1 | Q5SZD4 | 1.9646525 | 0.0448255 | GLYL3_HUMAN | |
| XM_013280258.1 | Q00630 | 1.9609111 | 0.0372242 | ICYB_MANSE | |
| XM_013287273.1 | P70169 | 1.7335803 | 0.0363557 | DOC2B_MOUSE | |
| XR_001225424.1 | B2S348 | 1.7255975 | 0.0036855 | SYI_TREPS | |
| XM_013294139.1 | Q7KML2 | 1.5104882 | 0.0043925 | ACOX1_DROME | |
| XM_013281121.1 | Q9UA35 | 1.4424482 | 0.000214 | S28A3_EPTST | |
| XM_013287297.1 | Q07075 | 1.4344746 | 0.014652 | AMPE_HUMAN | |
| XM_013278526.1 | P04694 | 1.4304328 | 4.22E-06 | ATTY_RAT | |
| XM_013281769.1 | Q93113 | 1.4082984 | 0.0004217 | GST1D_ANOGA | |
| XM_013284546.1 | Q9VZZ4 | 1.3242398 | 0.014652 | PXDN_DROME | |
| XM_013293212.1 | Q9BTE0 | 1.3146184 | 0.000499 | NAT9_HUMAN | |
| XM_013282652.1 | Q5XIM4 | 1.192563 | 0.0294286 | ATP5S_RAT | |
| XM_013291132.1 | Q9H0U6 | 1.170774 | 0.000214 | RM18_HUMAN | |
| XM_013292748.1 | Q9VJ26 | 1.1436627 | 0.0017239 | EFHD2_DROME | |

| | | | | |
|---|---|---|---|---|
| XM_013284680.1 | Q8NHP6 | 1.0543045 | 0.014652 | MSPD2_HUMAN |
| XM_013287288.1 | P30837 | 1.0397076 | 0.0159084 | AL1B1_HUMAN |
| XM_013285892.1 | Q9V6D6 | 1.0237154 | 0.0023031 | CP301_DROME |
| XM_013285172.1 | Q96LU5 | 1.018219 | 0.0003111 | IMP1L_HUMAN |
| XM_013292177.1 | O61577 | 1.015009 | 0.0443858 | KTNA1_STRPU |
| XM_013285904.1 | O95164 | 0.9989105 | 0.0248156 | UBL3_HUMAN |
| XM_013289770.1 | Q6PBY7 | 0.9527247 | 0.0130962 | TPC13_DANRE |
| XM_013280472.1 | Q9BV90 | 0.8990102 | 0.0256861 | SNR25_HUMAN |
| XM_013280974.1 | Q9UJK0 | 0.7170239 | 0.0485459 | TSR3_HUMAN |
| XM_013288199.1 | P06180 | 0.6992927 | 0.0428845 | HIBN_XENLA |
| XM_013284423.1 | O15498 | 0.6535519 | 0.029969 | YKT6_HUMAN |
| XM_013278936.1 | Q9U5V3 | 0.6147632 | 0.014652 | U239_DROSI |
| XM_013280038.1 | Q9NPB0 | 0.6024442 | 0.0280421 | SMDC1_HUMAN |
| XM_013294220.1 | B1MLL3 | 0.4877625 | 0.0463328 | GLGC_MYCA9 |
